# Multi-Omic Analysis of *Tyrophagus putrescentiae* Reveals Insights into the Allergen Complexity of Storage Mites

**DOI:** 10.1101/2022.10.21.513148

**Authors:** Angel Tsz-Yau Wan, Qing Xiong, Xiaojun Xiao, Kelvin Fu-Kiu Ao, Seok Woo Jang, Brian Shing-Hei Wong, Mingqiang Wang, Qin Cao, Cathy Sin-Hang Fung, Fook-Tim Chew, Baoqing Sun, Sai Ming Ngai, Ting-Fan Leung, Kyoung Yong Jeong, Xiaoyu Liu, Stephen Kwok-Wing Tsui

## Abstract

**Background:** The storage mite *Tyrophagus putrescentiae* is one of the major mites causing allergies in Chinese and Korean populations, but its allergen profile in incomplete when compared with that of house dust mites. Multiple genome-based methods have been introduced into the allergen study of mites and have enabled a better understanding of these medically important organisms.

**Objective:** We sought to reveal a comprehensive allergen profile of *Tyrophagus putrescentiae* and advance the allergen study of storage mites.

**Methods:** Based on a high-quality assembled and annotated genome, an *in silico* analysis was performed by searching reference sequences to identify allergens. Immunoassay ELISA assessed the allergenicities of recombinant proteins. MALDI-TOF mass spectrometry identified the IgE-binding proteins. Comparative genomics analysis was employed for the important allergen gene families.

**Results:** A complete allergen profile of *Tyrophagus putrescentiae* was revealed, including thirty-seven allergen groups (up to Tyr p 42). Among them, five novel allergens were verified using the sera of allergy patients. Massive allergen homologs were identified as the result of gene duplications in genome evolution. Proteomic identification again revealed a wide range of allergen homologs. In the NPC2 family and GSTs, comparative analysis shed light on the expansion and diversification of the allergen groups.

**Conclusion:** Using multi-omic approaches, the comprehensive allergen profile including massive homologs was disclosed in *Tyrophagus putrescentiae*, which revealed the allergen complexity of the storage mite and could ultimately facilitate the component-resolved diagnosis.

## Introduction

Astigmatic mites (suborder: Astigmata) comprise many medically important human household pests, and free-living species including house dust mites and storage mites are the major causes of human allergies ^1-6^. *Tyrophagus* (*T*.) *putrescentiae*, commonly referred to as the mold mite or cheese mite, is especially well-known as a storage mite that causes human allergic diseases such as atopic dermatitis, allergic rhinitis, and asthma ^7-10^. Unlike the house dust mites *Dermatophagoides* (*D*.) *farinae* and *D. pteronyssinus* mainly found in human household environments, *T. putrescentiae* has been reported to be in various environments including soil, hay, straw, and stored grain, as well as house dust ^7, 8^. Although *T. putrescentiae* is considered to be one of the major mites causing allergies in Chinese and Korean patients ^11-13^, it is less studied and has much fewer reported allergen groups than house dust mites ^14-16^. In addition, cross-reactivity has been frequently observed among astigmatic mites and causes difficulties in exploring *T. putrescentiae* allergies ^11, 17-19^.

In this study, we extensively analyzed the allergens of *T. putrescentiae* within the high-quality genome. Combining genomics, transcriptomics, and proteomics methods, multi-omic approaches were performed and a comprehensive allergen profile of *T. putrescentiae* was identified. Inspired by the identification of a range of allergen homologs, the comparative genomics analysis shed light on the evolution of allergen gene families. Taken together, our multi-omic analysis of *T. putrescentiae* allergens expanded our knowledge of storage mite allergies and revealed insights into the complex allergen profile of storage mites.

## Results

### Genome-wide analysis of allergen groups

Based on the genome assembly and annotation of *T. putrescentiae* ^1^, a genome-wide analysis was performed to explore the allergen groups. Using the mite allergens reported in the WHO/IUIS allergen nomenclature database as reference genes ^20^, thirty-seven allergen groups (up to Tyr p 42) were identified and composed a comprehensive allergen profile of *T. putrescentiae* (Table 1). To stay consistent with allergen group numbers of *D. farinae* and *D. pteronyssinus*, the previously reported Tyr p 34, 35 and 36 were renamed Tyr p 39, 41 and 42, respectively (Table 1). As reported, massive tandem gene duplications were identified in the genome of *T. putrescentiae* ^1^, which frequently caused gene fusion in the genome annotation. In addition, heterogeneity of the mite culture could cause artificial frameshifts in the genome annotation (Fig. E1). Therefore, all the allergen genes have been cautiously manually curated with high-coverage transcriptome data.

**Table 1.** Overview summary of all allergen genes identified in *T. putrescentiae*.

| Allergen ID | Isoallergen ID | Gene locus | Biochemical function | No. of exons | No. of amino acids | TPM value (Avg±Stdev) | homolog (Identity) |
| --- | --- | --- | --- | --- | --- | --- | --- |
| **Tyr p 1** | Tyr p 1.0101* | TP_010062.02 | Cysteine protease | 5 | 336 | 25±13 | ABM53753<br>Tyr p 1 (98.8%) |
|  | Tyr p 1.0201 | TP_003057.02 | Cysteine protease | 5 | 346 | 307±49 | ABM53753<br>Tyr p 1 (46.0%) |
|  | Tyr p 1.0301 | TP_003057.03 | Cysteine protease | 5 | 343 | 105±17 | ABM53753<br>Tyr p 1 (41.4%) |
|  | Tyr p 1.0401 | TP_003059.01 | Cysteine protease | 4 | 343 | 639±26 | ABM53753<br>Tyr p 1 (39.1%) |
|  | Tyr p 1.0501 | TP_011220.01 | Cysteine protease | 5 | 341 | 486±52 | ABM53753<br>Tyr p 1 (39.0%) |
| **Tyr p 2** | Tyr p 2.0101* | TP_003565.02 | NPC2 family | 2 | 141 | 11776±2386 | CAA73221<br>Tyr p 2 (99.3%) |
|  | Tyr p 2.0201* | TP_003567.01 | NPC2 family | 2 | 141 | 11776±2386 | CAA73221<br>Tyr p 2 (99.3%) |
|  | Tyr p 2.0301 | TP_019440.02 | NPC2 family | 3 | 142 | 192±37 | CAA73221<br>Tyr p 2 (70.4%) |
|  | Tyr p 2.0401 | TP_019491.01 | NPC2 family | 2 | 142 | 76±34 | CAA73221<br>Tyr p 2 (69.7%) |
|  | Tyr p 2.0501 | TP_003565.03 | NPC2 family | 2 | 144 | 1061±40 | CAA73221<br>Tyr p 2 (63.6%) |
|  | Tyr p 2.0601 | TP_003565.04 | NPC2 family | 2 | 142 | 6606±717 | CAA73221<br>Tyr p 2 (63.6%) |
| **Tyr p 3** | Tyr p 3.0101* | TP_012006.03 | Trypsin | 4 | 285 | 1374±691 | ABZ81991<br>Tyr p 3 (93.6%) |
|  | Tyr p 3.0201 | TP_012006.04 | Trypsin | 6 | 535 | 59±17 | ABZ81991<br>Tyr p 3 (69.8% / 60.7%) |
|  | Tyr p 3.0301 | TP_012006.02 | Trypsin | 4 | 288 | 4±3 | ABZ81991<br>Tyr p 3 (66.8%) |
|  | Tyr p 3.0401 | TP_014815.01 | Trypsin | 3 | 262 | 40218±11917 | ABZ81991<br>Tyr p 3 (51.5%) |
|  | Tyr p 3.0501 | TP_014833.01 | Trypsin | 3 | 262 | 40218±11917 | ABZ81991<br>Tyr p 3 (51.5%) |
|  | Tyr p 3.0601 | TP_022179.01 | Trypsin | 3 | 262 | 40218±11917 | ABZ81991<br>Tyr p 3 (51.5%) |
|  | Tyr p 3.0701 | TP_007838.01 | Trypsin | 3 | 286 | 7210±2050 | ABZ81991<br>Tyr p 3 (49.8%) |
| **Tyr p 4** | Tyr p 4.0101* | TP_001226.01 | Alpha-amylase | 5 | 518 | 1074±34 | ABZ81991<br>Tyr p 4 (99.0%) |
| **Tyr p 5** | Tyr p 5.0101 | TP_014187.01 | Unknown | 4 | 135 | 3590±407 | AAD10850<br>Blo t 5 (53.3%) |
|  | Tyr p 5.0201 | TP_014182.03 | Unknown | 2 | 128 | 691±276 | AAD10850<br>Blo t 5 (39.8%) |
|  | Tyr p 5.0301 | TP_014182.02 | Unknown | 2 | 236 | 11±3 | AAD10850<br>Blo t 5 (36.3%) |
| **Tyr p 6** | Tyr p 6.0101 | TP_021257.01 | Chymotrypsin | 4 | 275 | 23087±10331 | AAF28423<br>Der f 6 (55.4%) |
| **Tyr p 7** | Tyr p 7.0101* | TP_021945.01 | Bactericidal/permeability-increasing protein | 5 | 212 | 1119±193 | ABM53750<br>Tyr p 7 (99.1%) |
| **Tyr p 8** | Tyr p 8.0101 | TP_016261.02 | Glutathione S-transferase | 3 | 223 | 4990±109 | AGG10560<br>Tyr p 8 (63.9%) |
| **Tyr p 9** | Tyr p 9.0101 | TP_010332.01 | Serine protease | 3 | 270 | 43046±11849 | AAP57077<br>Der p 9 (55.1%) |
|  | Tyr p 9.0201 | TP_015698.02 | Serine protease | 2 | 299 | 300±53 | AAP57077<br>Der p 9 (48.7%) |
| **Tyr p 10** | Tyr p 10.0101* | TP_005966.01 | Tropomyosin | 8 | 284 | 36103±2786 | AAT40866<br>Tyr p 10 (98.6%) |
|  | Tyr p 10.0201 | TP_002355.01 | Tropomyosin | 7 | 253 | 3126±464 | AAT40866<br>Tyr p 10 (56.8%) |
| **Tyr p 11** | Tyr p 11.0101 | TP_012922.01 | Paramyosin | 12 | 839 | 6138±548 | AAM83103<br>Blo t 11 (91.5%) |
| **Tyr p 12** | Tyr p 12.0101 | TP_009564.03 | Chitin-binding domain containing protein | 2 | 167 | 1424±63 | AAA78904<br>Blo t 12 (49.2%) |
| **Tyr p 13** | Tyr p 13.0101* | TP_016572.02 | Fatty acid-binding protein | 2 | 131 | 34448±939 | AAU11502<br>Tyr p 13 (100%) |
|  | Tyr p 13.0201 | TP_016572.03 | Fatty acid-binding protein | 3 | 130 | 174146±22801 | AAU11502<br>Tyr p 13 (66.9%) |
|  | Tyr p 13.0301 | TP_005581.01 | Fatty acid-binding protein | 3 | 130 | 48327±13467 | AAU11502<br>Tyr p 13 (60.8%) |
|  | Tyr p 13.0401 | TP_013995.02 | Fatty acid-binding protein | 2 | 132 | 6136±745 | AAU11502<br>Tyr p 13 (52.8%) |
|  | Tyr p 13.0501 | TP_021718.01 | Fatty acid-binding protein | 2 | 132 | 4833±714 | AAU11502<br>Tyr p 13 (50.0%) |
| **Tyr p 14** | Tyr p 14.0101 | TP_017885.01 | Apolipoprotein | 6 | 1681 | 8650±255 | AAM21322<br>Der p 14 (45.5%) |
| **Tyr p 15** | Tyr p 15.0101 | TP_009564.02 | Chitinase | 4 | 560 | 1820±190 | AAAY84565<br>Der p 15 (72.5%) |
|  | Tyr p 15.0201 | TP_009563.02 | Chitinase | 4 | 554 | 2380±381 | AAAY84565<br>Der p 15 (72.0%) |
| **Tyr p 16** | Tyr p 16.0101 | TP_020865.01 | Gelsolin/villin | 8 | 497 | 963±33 | AAM64112<br>Der f 16 (54.5%) |
| **Tyr p 18** | Tyr p 18.0101 | TP_001639.01 | Chitinase | 4 | 464 | 4431±171 | AAM19082<br>Der f 18 (59.1%) |
|  | Tyr p 18.0201 | TP_021623.01 | Chitinase | 4 | 464 | 4431±171 | AAM19082<br>Der f 18 (59.1%) |
|  | Tyr p 18.0301 | TP_022773.02 | Chitinase | 4 | 466 | 1282±26 | AAM19082<br>Der f 18 (55.0%) |
| **Tyr p 20** | Tyr p 20.0101 | TP_001138.03 | Arginine kinase | 4 | 356 | 9868±886 | QOI58528<br>Tyr p 20 (88.2%) |
|  | Tyr p 20.0201 | TP_001138.02 | Arginine kinase | 5 | 392 | 17212±1384 | QOI58528<br>Tyr p 20 (87.1%) |
| **Tyr p 21** | Tyr p 21.0101 | TP_014185.01 | Unknown | 4 | 138 | 3472±202 | AAD10850<br>Blo t 21 (61.3%) |

**Table 1. Overview summary of all allergen genes in *T. putrescentiae***
| Allergen ID | Isoallergen ID | Gene locus | Biochemical function | No. of exons | No. of amino acids | TPM value (Avg±Stdev) | homolog (Identity) |
| --- | --- | --- | --- | --- | --- | --- | --- |
| **Tyr p 23** | Tyr p 23.0101 | TP_019120.02 | Peritrophin-like protein | 1 | 229 | 438±25 | ACB46292<br>Der p 23 (45.1%) |
| **Tyr p 24** | Tyr p 24.0101 | TP_009264.02 | UQCRB protein | 4 | 118 | 47455±5790 | AGI78542<br>Der f 24 (63.6%) |
| **Tyr p 25** | Tyr p 25.0101 | TP_010799.02 | Triosphosphate isomerase | 4 | 247 | 15900±1082 | AGC56216<br>Der f 25 (79.4%) |
| **Tyr p 26** | Tyr p 26.0101 | TP_009336.02 | Myosin light-chain | 5 | 158 | 49121±1725 | QAT18638<br>Der p 26 (87.8%) |
| **Tyr p 28** | Tyr p 28.0101 | TP_011342.02 | Heat shock protein 70 | 3 | 659 | 14066±3095 | AOD75395<br>Tyr p 28 (90.9%) |
|  | Tyr p 28.0201 | TP_011350.01 | Heat shock protein 70 | 3 | 655 | 5912±792 | AOD75395<br>Tyr p 28 (90.6%) |
|  | Tyr p 28.0301 | TP_022700.02 | Heat shock protein 70 | 5 | 644 | 1±2 | AOD75395<br>Tyr p 28 (83.2%) |
|  | Tyr p 28.0401 | TP_015983.01 | Heat shock protein 70 | 1 | 645 | 1817±36 | AOD75395<br>Tyr p 28 (80.8%) |
|  | Tyr p 28.0501 | TP_000248.01 | Heat shock protein 70 | 2 | 639 | 1322±107 | AOD75395<br>Tyr p 28 (78.7%) |
| **Tyr p 29** | Tyr p 29.0101 | TP_004508.01 | Cyclophilin | 3 | 331 | 315±121 | QAT18640<br>Der p 29 (69.3%) |
| **Tyr p 30** | Tyr p 30.0101 | TP_016485.02 | Ferritin | 1 | 107 | 1015±192 | QAT18641<br>Der p 30 (83.5%) |
|  | Tyr p 30.0201 | TP_021342.02 | Ferritin | 3 | 175 | 51401±3389 | QAT18641<br>Der p 30 (83.5%) |
|  | Tyr p 30.0301 | TP_021345.02 | Ferritin | 2 | 347 | 1090±179 | QAT18641<br>Der p 30 (83.5%) |
|  | Tyr p 30.0401 | TP_021341.01 | Ferritin | 2 | 172 | 13121±440 | QAT18641<br>Der p 30 (81.6%) |
| **Tyr p 31** | Tyr p 31.0101 | TP_010684.02 | Cofilin | 4 | 148 | 49319±7596 | QAT18642<br>Der p 31 (89.2%) |
| **Tyr p 32** | Tyr p 32.0101 | TP_016449.02 | Inorganic pyrophosphatase | 2 | 294 | 5123±879 | QAT18643<br>Der p 32 (70.0%) |
| **Tyr p 33** | Tyr p 33.0101 | TP_014283.02 | Alpha-tubulin | 6 | 451 | 4882±1325 | AIO08861<br>Der f 33 (83.7%) |
|  | Tyr p 33.0201 | TP_000522.01 | Alpha-tubulin | 6 | 450 | 22499±4863 | AIO08861<br>Der f 33 (82.7%) |
|  | Tyr p 33.0301 | TP_000118.02 | Alpha-tubulin | 8 | 347 | 930±110 | AIO08861<br>Der f 33 (80.2%) |
|  | Tyr p 33.0401 | TP_019973.02 | Alpha-tubulin | 6 | 446 | 4868±1279 | AIO08861<br>Der f 33 (79.6%) |
|  | Tyr p 33.0501 | TP_008758.02 | Alpha-tubulin | 7 | 450 | 5301±1043 | AIO08861<br>Der f 33 (78.5%) |
| **Tyr p 34** | Tyr p 34.0101 | TP_020961.02 | Rid-like protein | 1 | 137 | 4617±121 | BAV90601<br>Der f 34 (59.1%) |
|  | Tyr p 34.0201 | TP_012005.01 | Rid-like protein | 1 | 138 | 777±70 | BAV90601<br>Der f 34 (37.1%) |
| **Tyr p 36** | Tyr p 36.0101 | TP_004901.01 | Unknown | 2 | 223 | 5679±33 | ATI08932<br>Der f 36 (52.4%) |
|  | Tyr p 36.0201 | TP_006471.02 | Unknown | 3 | 222 | 1122±85 | ATI08932<br>Der f 36 (48.9%) |
|  | Tyr p 36.0301 | TP_020584.01 | Unknown | 3 | 235 | 9155±619 | ATI08932<br>Der f 36 (47.1%) |
|  | Tyr p 36.0401 | TP_010549.02 | Unknown | 3 | 236 | 97±20 | ATI08932<br>Der f 36 (41.7%) |
| **Tyr p 37** | Tyr p 37.0101 | TP_000195.01 | Chitin binding protein | 3 | 248 | 1716±221 | QBF67839<br>Der f 37 (44.9%) |
|  | Tyr p 37.0201 | TP_019138.01 | Chitin binding protein | 2 | 218 | 720±28 | QBF67839<br>Der f 37 (44.5%) |
| **Tyr p 38** | Tyr p 38.0101 | TP_012405.02 | Bacteriolytic enzyme | 4 | 153 | 819±340 | QHQ72282<br>Der f 38 (58.8%) |
|  | Tyr p 38.0201 | TP_012405.03 | Bacteriolytic enzyme | 3 | 153 | 22±5 | QHQ72282<br>Der f 38 (54.2%) |
|  | Tyr p 38.0301 | TP_019308.02 | Bacteriolytic enzyme | 2 | 144 | 47±10 | QHQ72282<br>Der f 38 (50.7%) |
| **Tyr p 39** | Tyr p 39.0101* | TP_005620.02 | Troponin C | 5 | 153 | 46444±821 | ACL36923<br>Tyr p 34# (100%) |
| **Tyr p 40** | Tyr p 40.0101 | TP_011775.02 | Thioredoxin-like protein | 3 | 165 | 28168±2971 | UJQ69787<br>Der f 40 (68.6%) |
| **Tyr p 41** | Tyr p 41.0101 | TP_014714.03 | Aldehyde dehydrogenase | 5 | 487 | 10169±1661 | AOD75396<br>Tyr p 35# (85.1%) |
|  | Tyr p 41.0201 | TP_014714.02 | Aldehyde dehydrogenase | 5 | 481 | 18053±266 | AOD75396<br>Tyr p 35# (74.7%) |
|  | Tyr p 41.0301 | TP_004844.03 | Aldehyde dehydrogenase | 5 | 483 | 138±27 | AOD75396<br>Tyr p 35# (67.8%) |
|  | Tyr p 41.0401 | TP_004844.02 | Aldehyde dehydrogenase | 5 | 483 | 2220±78 | AOD75396<br>Tyr p 35# (67.6%) |
| **Tyr p 42** | Tyr p 42.0101* | TP_007980.01 | Profilin | 3 | 131 | 61340±10960 | AOD75399<br>Tyr p 36# (91.6%) |

\* These genes were suggested as the identical allergen genes by the high identity to those reported in the WHO/IUIS Allergen Nomenclature database.

A wide range of gene expansion was observed in the allergen gene families and only those top matched homologs (or isoallergens) were listed, such as five homologs of the cysteine protease Tyr p 1 (Table 1). In our naming rules, we used the first and last two digits after the decimal point to differentiate the different genes and isoforms, respectively. For example, the two homologs of Tyr p 2, TP_003565.02 and TP_003567.01, shared identical protein and coding sequences but were named Tyr p 2.0101 and Tyr p 2.0201, respectively (Table 1). Multiple isoforms originating from the same gene locus were not included in this study, such as those caused by single-nucleotide polymorphisms or alternative splicing.

The gene expression levels were analyzed using two transcriptomic datasets of adult *T. putrescentiae* mites. Unlike the allergen gene expression of *D. farinae* and *D. pteronyssinus* ^21^, the group 1 allergens (cysteine proteases) of *T. putrescentiae* were expressed at extremely low levels, while the homolog of group 13 allergen, Tyr p 13.0201, exhibited the highest expression level (Fig. 1A). The gene synteny alignment revealed that many homologs were tandemly arrayed (Fig. 1B), which was the result of tandem gene duplication ^1^. In the group 1 allergen, the best matched gene Tyr p 1.0101 (gene locus: TP_010062.02) shared as high as 98.8% identity with the reported Tyr p 1 (Table 1). Three homologs (Tyr p 1.0201, Tyr p 1.0301 and Tyr p 1.0401) were tandemly arrayed and located closely (less than 5 kb in distance) on the same strand (Fig. 1B).

**Fig. 1.**
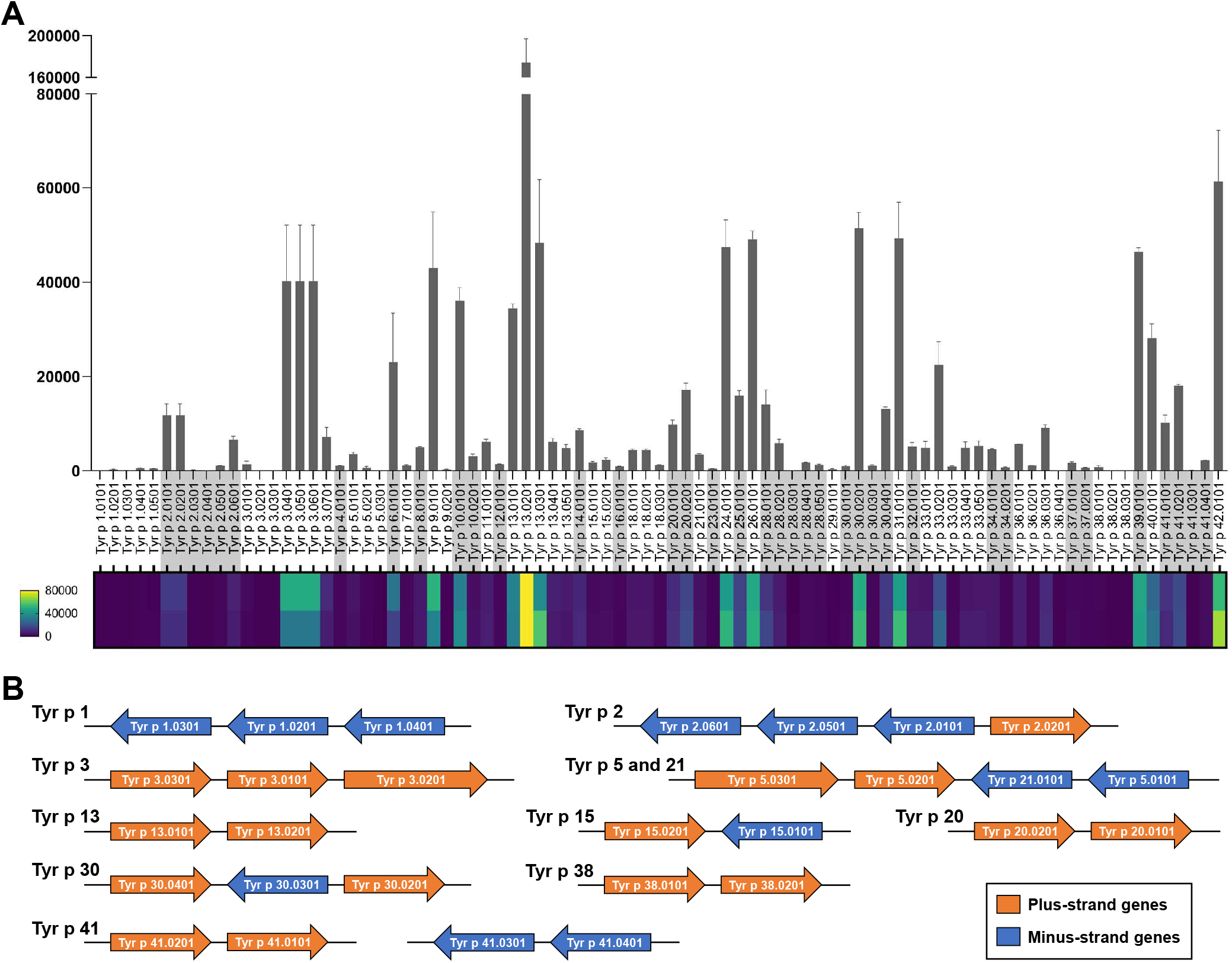
Allergen expression levels and tandem gene synteny alignment. **(A)** The expression levels of all identified allergens in *T. putrescentiae* (Table 1) were quantified as transcripts per million (TPM) using two transcriptome data of adult mites (TP1 and TP3 of SRR13837414). The error bars represent standard deviation in the bar chart. The maximum value was set as 80000 in the lower heatmap. **(B)** Gene synteny alignment of tandemly arrayed allergen genes. All the distances between neighboring genes are below 5 Kb.

Group 2 allergens belong to the Niemann-Pick protein type C2 (NPC2) family and are considered the major allergens of storage mites including *T. putrescentiae* ^22, 23^. In the genome of *T. putrescentiae*, we identified up to six homologs of Tyr p 2, including the two homologous genes (Tyr p 2.0101 and Tyr p 2.0201) shared identical sequences, and as high as 99.3% identity with the reported Tyr p 2 (GenBank accession: CAA73221, Table 1). Tyr p 2.0101 and Tyr p 2.0201 shared identical protein and coding sequences and the transcriptomic reads could not differentiate two identical genes. Therefore, Tyr p 2.0101 and Tyr p 2.0201 shared the same expression level (Table 1). Similarly, four homologs of Tyr p 2 (Tyr p 2.0101, Tyr p 2.0201, Tyr p 2.0501 and Tyr p 2.0601) were tandemly arrayed and Tyr p 2.0201 was located on the opposite strand (Fig. 1B). Considering that group 22 and 35 allergens were both characterized as members of the NPC2 family ^24-26^, we will perform a comparative analysis in astigmatic mites to further explore this important gene family.

Group 3, 6 and 9 allergens are all serine proteases that have been reported to have massive tandem gene duplication in astigmatic mites ^1^. The best matched Tyr p 3.0101 shared 93.6% identity with the reported Tyr p 3 (Table 1, Fig. E2A). Of the seven homologs of Tyr p 3, three genes (Tyr p 3.0101, Tyr p 3.0201 and Tyr p 3.0301) were tandemly arrayed (Fig. 1B) and Tyr p 3.0201 was identified as a dimeric trypsin containing two duplicated domains, while Tyr p 3.0401, Tyr p 3.0501 and Tyr p 3.0601 shared identical protein and coding sequences. Consistent with a previous report, the global expression levels of serine protease allergens were relatively higher than those of group 1 cysteine proteases ^27^.

Only one alpha-amylase gene of *T. putrescentiae* ^1, 28^, Tyr p 4.0101, was identified as highly similar to the reported Tyr p 4 (99.0% identity, Table 1). Group 5 and 21 allergens are the major allergens of *Blomia* (*B*.) *tropicalis* ^29, 30^, but have not been reported in *T. putrescentiae*. We identified three homologs of Tyr p 5 (including dimeric Tyr p 5.0301) and one of Tyr p 21 (Table 1), all of which were tandemly arrayed (Fig. 1B).

Tyr p 7.0101 shared 99.1% identity with the reported Tyr p 7 (Table 1), while unexpectedly, the best matched homolog of the group 8 allergen Tyr p 8.0101 shared only 63.9% identity with the reported Tyr p 8 (GenBank accession: AGG10560, Fig. E2B). Two tandemly arrayed homologs of Tyr p 20, Tyr p 20.0101 and Tyr p 20.0201 (gene locus: TP_001138.03 and TP_001138.02) shared 88.2% and 87.1% identities with the reported Tyr p 20 (GenBank accession: QOI58528, Fig. E2C), respectively. In addition, the best matched genes of Tyr p 41 and 42 shared 85.1% and 91.6% identities with the reported allergens, respectively (Fig. E2D and E). Due to space limitations, we will not discuss other allergen groups exhaustively.

To evaluate the allergenicity of the newly identified allergens (Table 1), recombinant proteins were cloned and expressed, and the IgE reactivity was assessed by the enzyme-linked immunosorbent assay (ELISA) with the sera of *T. putrescentiae* sensitized patients (Table E1). The recombinant proteins showed various allergenicities (Fig. E3). For the two serine protease allergens, the recombinant proteins rTyr p 6.0101 and rTyr p 9.0101 presented low allergenicities of 11.1% and 22.2%, respectively (Fig. E3A and B). Similarly, the chitinase rTyr p 18.0101 tested positive in only two samples (11.1%, Fig. E3C). For the arginine kinase and the myosin light-chain, rTyr p 20.0101 and rTyr p 26.0101 exhibited higher allergenicities, 44.4% and 50.0%, respectively (Fig. E3D and E). Although all the patients were diagnosed as *T. putrescentiae* sensitized, only 11/18 (61.1%) of the serum samples showed positive responses to the crude protein extract (Fig. E3F).

### Proteomic identification of IgE-binding proteins

In addition to the *in silico* analysis, proteomic identification was performed using the pooled sera of allergy patients to identify *T. putrescentiae* allergens (Table E2). In total, thirty-one protein spots of *T. putrescentiae* bound by IgE in the serum samples underwent *de novo* peptide sequencing by MALDI-TOF mass spectrometry (Fig. E4), and then the sequenced peptides were searched against the annotated protein sequences of the *T. putrescentiae* genome.

A range of our identified allergens could be found in the spots, including members of Tyr p 1, 2, 3, 8, 10, 13, 20, 21, 25, 28 and 39 (Table 2). Among them, the best matched homologs in the spots included Tyr p 2.0101, Tyr p 10.0101, Tyr p 20.0101, Tyr p 21.0101, Tyr p 25.0101, Tyr p 28.0101 and Tyr p 39.0101. In particular, Tyr p 21.0101 was matched by four spots, while Tyr p 10.0101, Tyr p 20.0101 and Tyr p 25.0101 were each found in two spots (Table 2). Some genes were suggested as allergen homologs but shared relatively low identity with the reported allergens, so they were not listed in Table 1 and labeled ungroup allergens (Table 2). However, in Tyr p 2, Tyr p 2.0101/Tyr p 2.0201, Tyr p 2.0601 and an ungrouped Tyr p 2 (gene locus: TP_020235.01) were matched by the sequenced peptides (Table 2). For Tyr p 28, Tyr p 28.0101, Tyr p 28.0201 and an ungrouped Tyr p 28 (gene locus: TP_006940.01) were identified in the spots (Table 2).

**Table 2.** MALDI-TOF MS results of IgE-bound *T. putrescentiae* proteins.

| ID on 2D gel | Accession | Allergen | Protein | Score | MW (kDa) | pI | No. of peptides | Coverage (%) |
| --- | --- | --- | --- | --- | --- | --- | --- | --- |
| 1 | TP_005966.01 | Tyr p 10.0101 | Tropomyosin | 334.36 | 33 | 4.78 | 6 | 21.8 |
| 2 | TP_005966.01 | Tyr p 10.0101 | Tropomyosin | 265.73 | 33 | 4.78 | 4 | 14.8 |
| 6 | TP_010799.01 | Tyr p 25.0101 | Triosphosphate isomerase | 674.93 | 26.8 | 6.13 | 6 | 41.3 |
| 8 | TP_010799.01 | Tyr p 25.0101 | Triosphosphate isomerase | 610.06 | 26.8 | 6.13 | 6 | 41.3 |
|  | TP_007838.01 | Tyr p 3.0701 | Trypsin | 57.33 | 29.9 | 7.1 | 2 | 7.7 |
| 9 | TP_004226.01 | -- | Sod1 Superoxide dismutase [Cu-Zn] | 576.51 | 15.6 | 5.92 | 5 | 51 |
| 10 | TP_003565.04 | Tyr p 2.0601 | NPC2 family | 51.72 | 15.1 | 6.99 | 1 | 7.7 |
| 11 | TP_003565.04 | Tyr p 2.0601 | NPC2 family | 354.79 | 15.1 | 6.99 | 4 | 41.5 |
|  | TP_004226.01 | -- | Sod1 Superoxide dismutase [Cu-Zn] | 115.71 | 15.6 | 5.92 | 3 | 23.2 |
|  | TP_014856.01 | -- | Unknown | 42.84 | 24.9 | 6.93 | 1 | 6.2 |
| 13 | TP_020235.01 | Ungrouped Tyr p 2 *<br>(44.0%, CAA73221) | NPC2 family | 60.89 | 15.6 | 7.1 | 2 | 17.1 |
| 15 | TP_004518.01 | -- | FK506-binding protein | 296.13 | 11.5 | 6.72 | 4 | 40.7 |
|  | TP_014185.01 | Tyr p 21.0101 | Unknown | 180.4 | 15.3 | 7.83 | 3 | 30.4 |
| 16 | TP_014185.01 | Tyr p 21.0101 | Unknown | 538.38 | 15.3 | 7.83 | 6 | 46.4 |
| 17 | TP_011560.01 | Ungrouped Tyr p 13 *<br>(37.4%, AAU11502) | Fatty acid-binding protein | 392.65 | 17.5 | 8.61 | 5 | 42.9 |
|  | TP_003565.02/<br>TP_003567.01 | Tyr p 2.0101/<br>Tyr p 2.0201 | NPC2 family | 184.57 | 14.8 | 8.51 | 2 | 27.7 |
| 18 | TP_014185.01 | Tyr p 21.0101 | Unknown | 161.6 | 15.3 | 7.83 | 3 | 30.2 |
| 19 | TP_014185.01 | Tyr p 21.0101 | Unknown | 46.76 | 15.3 | 7.83 | 1 | 10.1 |
| 20 | TP_004953.01 | -- | Unknown | 132.75 | 34.2 | 9.96 | 3 | 17.9 |
|  | TP_005006.01 | -- | Unknown | 61.66 | 34.2 | 6.9 | 2 | 11.5 |
| 23 | TP_011350.01 | Tyr p 28.0201 | Heat shock protein 70 | 267.96 | 71.5 | 5.29 | 4 | 8.1 |
|  | TP_011342.01 | Tyr p 28.0101 | Heat shock protein 70 | 141.09 | 122 | 5.33 | 3 | 3.2 |
|  | TP_006940.01 | Ungrouped Tyr p 28 *<br>(51.3%, AOD75395) | Heat shock protein 70 | 65.45 | 74.4 | 5.64 | 2 | 3.8 |
| 24 | TP_001283.01 | -- | Nucleoside diphosphate kinase A1 | 280.29 | 23.3 | 8.32 | 5 | 24.2 |
|  | TP_006307.01 | -- | HEXBP DNA-binding protein HEXBP | 173.74 | 14.1 | 8.26 | 3 | 40.8 |
|  | TP_020843.01 | -- | CYPA Peptidyl-prolyl cis-trans isomerase | 81.44 | 28.8 | 10.08 | 1 | 5.3 |
|  | TP_005620.02 | Tyr p 39.0101 | Troponin C | 52.78 | 17.7 | 4.04 | 1 | 9.8 |
| 25 | TP_001138.03 | Tyr p 20.0101 | Arginine kinase | 421.42 | 40.1 | 6.23 | 5 | 21.7 |
|  | TP_009769.01 | -- | FBPA Fructose-bisphosphate aldolase | 84.84 | 38.7 | 6.55 | 3 | 14.3 |
|  | TP_008599.01 | Ungrouped Tyr p 1 *<br>(28.2%, ABM53753) | Cysteine protease | 69.95 | 39.5 | 5.9 | 2 | 10 |
| 26 | TP_001138.03 | Tyr p 20.0101 | Arginine kinase | 427.58 | 40.1 | 6.23 | 5 | 21.7 |
|  | TP_009769.01 | -- | FBPA Fructose-bisphosphate aldolase | 71.34 | 38.7 | 6.55 | 2 | 6.6 |
| 29 | TP_007523.01 | -- | pdi-2 Protein disulfide-isomerase 2 | 236.42 | 54.9 | 4.95 | 4 | 10.5 |
|  | TP_006736.01 | -- | ATPsynbeta ATP synthase subunit beta, mitochondrial | 99.41 | 56.1 | 5.13 | 3 | 8.4 |
| 31 | TP_013474.01 | -- | Gpx5 Epididymal secretory glutathione peroxidase | 139.51 | 25.8 | 7.12 | 2 | 9.7 |
|  | TP_014174.01 | Ungrouped Tyr p 8 *<br>(50.7%, AGG10560) | Glutathione S-transferase | 103.13 | 24.9 | 8.95 | 2 | 12.7 |
|  | TP_019125.01 | -- | Cuticle protein 16.8 | 87.72 | 25.6 | 8.92 | 1 | 11.6 |

\* These genes were suggested as allergen homologs but shared relatively low identity with the reported allergens, so that they were not listed in Table 1. The identity percentage and the GenBank accession of the reference allergen were noted in the bracket.

The ungrouped Tyr p 1 (gene locus: TP_008599.01) was identified and shared only 28.2% identity with the reported Tyr p 1 (Table 2). This cysteine protease homolog (gene locus: TP_008599.01) was estimated to be expressed over two hundred times more than the best matched Tyr p 1.0101 (Table 1) and lower than 50% of that of Tyr p 3.0401 (Fig. 1A). Similarly in Tyr p 13, the ungrouped homolog (gene locus: TP_011560.01) was expressed at over 50% higher levels than the highly expressed Tyr p 13.0201. Therefore, we proposed that the gene expression level was a crucial factor in proteomic identification. In addition, an ungrouped homolog of Tyr p 8 (gene locus: TP_014174.01) was identified, but not the *in silico* predicted Tyr p 8.0101 (Table 1). Comparative analysis of glutathione S-transferases (GSTs) will be performed to further explore group 8 allergens.

Other proteins such as the FK506-binding protein (gene locus: TP_004518.01) were identified in one spot (Table 2). The FK506-binding protein was expressed, and its allergenicity was assessed to be 22.2% (Fig. E5). Notably, the serum samples that we used were assessed as positive for *D. pteronyssinus* (Table S1) but have not been tested for *T. putrescentiae*.

### Comparative analysis of the NPC2 family and GSTs

Comparative genomic analysis has revealed the divergent evolution of astigmatic mites ^1^. Our multiomic identification of allergen homologs in *T. putrescentiae* indicated the necessity of comparative analysis of the allergen gene families.

Group 2 allergens belonging to the NPC2 family are especially responsible for the cross-reactivity among mite species ^31, 32^. To explore the NPC2 family, we collected the genes of six astigmatic mites and performed phylogenetic analysis with the reported group 2 allergens. In the phylogenetic tree, the NPC2 family genes were divided into eight clusters, i.e., N1-3 and C1-5 (Fig. 2A). All reported allergen genes were in Cluster N1-3, while none of the genes in Cluster C1-5 were reported to be allergens. Cluster N1 contained Blo t 2, Tyr p 2, Lep d 2 and Gly d 2 and Der f 35, as well as Pso o 2 (UniProt ID: Q965E2) of *Psoroptes* (*P*.) *ovis*. Cluster N2 covered group 2 allergens of house dust mites including Der f 2, Der p 2 and Eur m 2, while Cluster N3 contained Der f 22. The gene synteny alignment suggested that the NPC2 gene of *S. scabiei* in N1 decayed, while that of *B. tropicalis* was tandemly duplicated (Fig. 2B). Cluster N2 is proposed to be unique in psoroptid mites, but decayed in *P. ovis*, according to the gene synteny alignment (Fig. 2C), while a 38-aa insertion was identified in the N-terminus of SS_011027.01 in *S. scabiei* (Fig. E6). At the gene expression level, most highly expressed genes were in Cluster N1-3 (Fig. E7), except one *S. scabiei* gene in Cluster C1, and one *D. farinae* gene and one *T. putrescentiae* gene in Cluster C3.

**Fig. 2.**
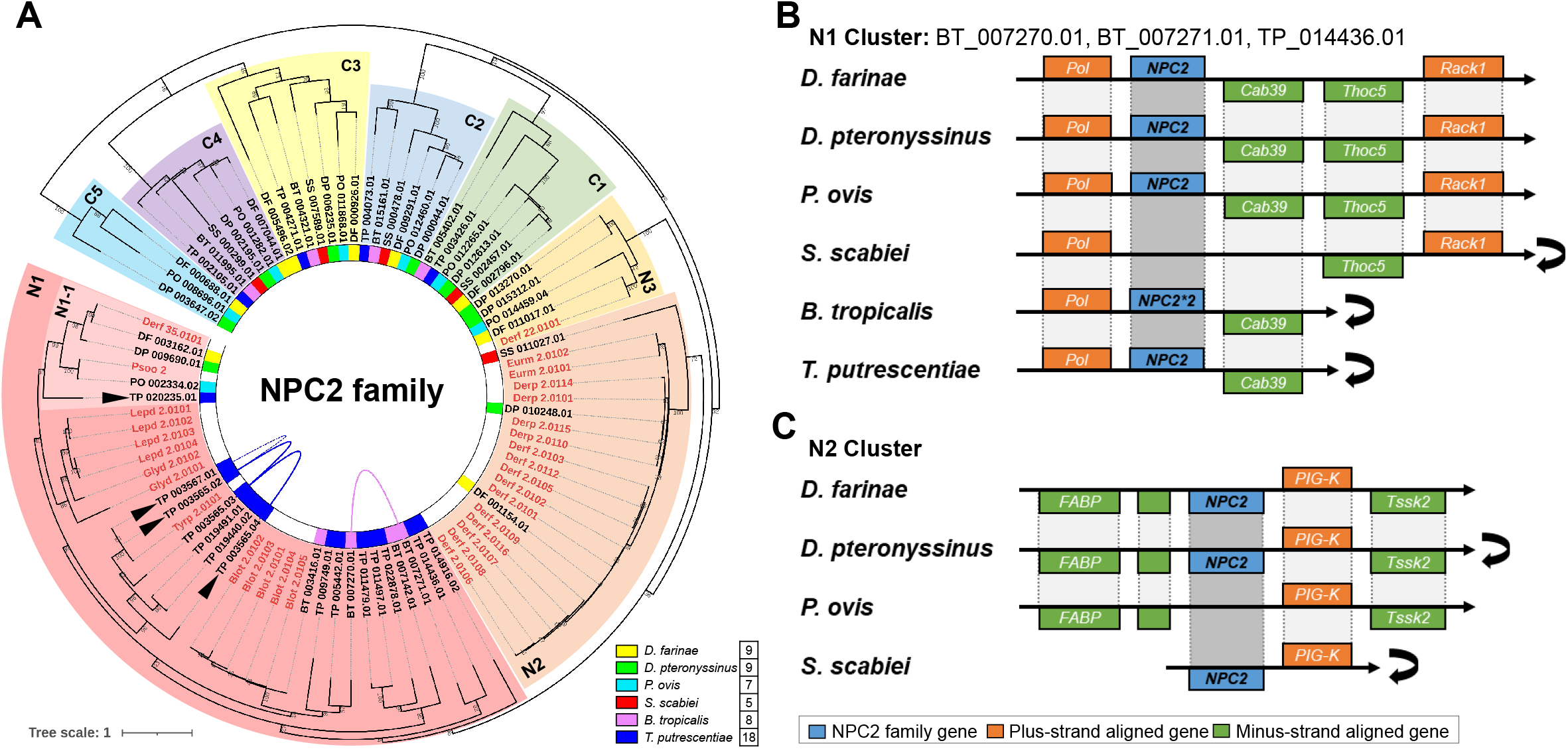
Comparative analysis of NPC2 family. **(A)** Phylogenetic analysis of NPC2 family of six astigmatic mites, including two house dust mites, *Dermatophagoides* (*D*.) *farinae* and *D. pteronyssinus*, two parasitic mites, *Psoroptes* (*P*.) *ovis* and *Sarcoptes* (*S*.) *scabiei*, and two canonical storage mites, *Blomia* (*B*.) *tropicalis* and *T. putrescentiae*. The NPC2 genes were divided into 8 clusters, i.e., N1-3 and C1-5. The NPC2 allergens (highlighted in red) include group 2, 22 and 35 of astigmatic mites collected from the WHO/IUIS nomenclature database and Pso o 2 of *P. ovis* from UniProt database (ID: Q965E2). Tandemly arrayed genes and proximally arrayed genes (separated by no more than 10 genes) were connected by curved solid lines and dotted lines, respectively. Four black triangles marked the four NPC2 genes identified in MS (Table 2). **(B)** Gene synteny alignment of N1 Cluster. The NPC2 gene was not found in *S. scabiei*, but tandemly duplicated in *B. tropicalis*. **(C)** Gene synteny alignment of N2 Cluster. No homolog was found in *P. ovis*. The black turnover arrow means after reverse complement.

In the Mu-class GST Mu, group 8 allergen can also cause cross-reactivity among astigmatic mites ^33^. Similarly, the GSTs of six astigmatic mites were phylogenetically analyzed (Fig. E8A). Six clusters were divided, and all reported allergens were in Cluster GST Mu. Unexpectedly, none of the GST genes were identical to the reported Tyr p 8 (GenBank accession: AGG10560). Intriguingly, massive tandem gene duplications were observed especially in the allergen cluster, Cluster GST Mu (Fig. E8A). Unlike the ungrouped Tyr p 1 and 13 (Table 2), the proteomically identified GST (ungrouped Tyr p 8, gene locus: TP_014174.01) was a lowly expressed homolog, which was unexpected (Fig. E8B).

## Discussion

In the omics era of mite study, multiple genome-based approaches have been boosting our understanding of these medically important organisms and benefitting the component-resolved diagnosis of mite allergies ^1, 21, 34, 35^. Based on the high-quality genome, we performed multi-omic analysis to explore the allergens of the storage mite *T. putrescentiae*. A comprehensive allergen profile of *T. putrescentiae* was identified using multi-omic approaches, and a wide range of allergen homologs were observed in both *in silico* and proteomic analyses. The comparative analysis of the NPC2 family and GSTs provided an evolutionary perspective of the allergen groups that are considered important for cross-reactivity. Taken together, this integrative analysis of *T. putrescentiae* allergens provided a systematic understanding of the allergen complexity of storage mites.

With the aid of high-quality genome and transcriptome data, our *in silico* analysis revealed a comprehensive allergen profile, including thirty-seven allergen groups (up to Tyr p 42, Table 1). All the reported allergens of *T. putrescentiae* could be found in the list, except Tyr p 8, 20 and 41 which lack highly similar (>90% identity) homologs. The discrepancy can be attributed to intragenic divergence of *T. putrescentiae*, except that the best match of Tyr p 8 shared only 63.9% identity. The gene expression profile of *T. putrescentiae* allergens was distinct from that of the house dust mites *D. farinae* and *D. pteronyssinus*, in which the major house dust mite allergen, group 1 allergens were expressed at extremely low levels (Fig. 1A). Gene family expansion was frequently observed in these allergen groups and resulted in a wide range of allergen homologs, in which massive tandem gene duplications were identified such as in group 1 and 2 (Fig. 1B). These homologs contribute to the high diversity of *T. putrescentiae* allergens. To evaluate the IgE-binding reactivity of novel allergens, recombinant proteins were cloned, expressed, and assessed by ELISA with *T. putrescentiae*-sensitized patient sera. Despite low allergenicities, five proteins, Tyr p 6.0101, Tyr p 9.0101, Tyr p 18.0101, Tyr p 20.0101 and Tyr p 26.0101, were suggested to be novel allergens of *T. putrescentiae* by ELISA experiments (Fig. E3A-E).

Apart from *in silico* analysis, proteomic identification using pooled allergy patient sera provided more interesting information for *T. putrescentiae* allergens. Except for the reported allergens, novel allergens such as Tyr p 21.0101 were well matched by the sequenced peptides of the protein spots bound to IgE (Table 2). Moreover, a wide range of allergen homologs appeared in the proteomic identification, including ungrouped homologs such as Tyr p 1, 2 and 8 (Table 2). In addition, other proteins that have never been reported as allergens could also be found, such as the FK506-binding protein (gene locus: TP_004518.01) that was suggested as a novel allergen by ELISA (Fig. E5). The pooled sera used in proteomic identification were assessed as positive for *D. pteronyssinus* but unknown for *T. putrescentiae*, which may be involved in the cross-reactivity problem at the beginning. The proteomic identification revealed complex allergen homologs, which is concurrent with the *in silico* analysis (Table 1).

Not only the *in silico* analysis but also the proteomic identification revealed massive Tyr p 2 homologs in the NPC2 family (Table 1 and 2). Group 2 allergens contribute to the cross-reactivity among mite species ^31, 32^. Comparative analysis revealed the evolution of the important NPC2 family of astigmatic mites and the relationship among group 2, 22 and 35 allergens. (Fig. 2) Group 22 allergens were quite unique in *D. farinae, D. pteronyssinus* and the sheep scab mite *P. ovis*, while group 35 allergen of *D. farinae*, Der f 35, were clustered with the diversified NPC2 genes of storage mites including the group allergens Tyr p 2 and Lep d 2 (Fig. 2A). Additionally, we explored the unusual group 8 allergens (characterized as GSTs) and observed frequent tandem gene duplications especially in the cluster GST mu covering the group 8 allergens of mites (Fig. E8). The reported Tyr p 8 (GenBank accession: AGG10560) could not find a highly similar homolog. Unexpectedly, the proteomically identified Tyr p 8 homolog was expressed at a low level (gene locus: TP_014174.01). Using the NPC2 family and GSTs as examples, our comparative analysis sheds light on the diversification of allergen gene families in mite species.

## Abbreviations used

ELISA: enzyme-linked immunosorbent assay
GST: glutathione S-transferases
MALDI-TOF: matrix-assisted laser desorption/ionization-time of flight
NPC2: Niemann-Pick protein type C2

## Acknowledgments

We would like to thank Prof. Yubao CUI of the Affiliated Wuxi People’s Hospital of Nanjing Medical University for providing the *T. putrescentiae* mites and the staff members in the Shenzhen Key Laboratory of Allergy for the mite culturing work.

## Data availability

The genome and sequencing data of *T. putrescentiae* are deposited in NCBI database under BioProject accession PRJNA706095. The *in silico* identified allergen sequences were uploaded in NCBI GenBank database under accessions OP558975-OP559059.

## References

1. Xiong Q, Wan AT-Y, Liu X, et al. Comparative Genomics Reveals Insights into the Divergent Evolution of Astigmatic Mites and Household Pest Adaptations. Molecular Biology and Evolution. 2022;39(5) doi:10.1093/molbev/msac097

2. Klimov PB, Oconnor B. Is Permanent Parasitism Reversible?—Critical Evidence from Early Evolution of House Dust Mites. Systematic Biology. 2013;62(3):411–423. doi:10.1093/sysbio/syt008

3. Domes K, Althammer M, Norton RA, Scheu S, Maraun M. The phylogenetic relationship between Astigmata and Oribatida (Acari) as indicated by molecular markers. Experimental & applied acarology. 2007;42(3):159–71. doi:10.1007/s10493-007-9088-8

4. Norton RA. Morphological evidence for the evolutionary origin of Astigmata (Acari: Acariformes). Experimental & applied acarology. 1998;22(10):559–594.

5. Sakata T, Norton RA. Opisthonotal gland chemistry of early-derivative oribatid mites (Acari) and its relevance to systematic relationships of Astigmata. International Journal of Acarology. 2001;27(4):281–292.

6. Dabert M, Witalinski W, Kazmierski A, Olszanowski Z, Dabert J. Molecular phylogeny of acariform mites (Acari, Arachnida): Strong conflict between phylogenetic signal and long-branch attraction artifacts. Molecular Phylogenetics and Evolution. 2010/07/01/ 2010;56(1):222–241. doi:https://doi.org/10.1016/j.ympev.2009.12.020

7. Green WF, Woolcock AJ. Tyrophagus putrescentiae: an allergenically important mite. Clinical Experimental Allergy. 1978;8(2):135–144. doi:10.1111/j.1365-2222.1978.tb00458.x

8. Arlian LG, Geis DP, Vyszenski-Moher DL, Bernstein IL, Gallagher JS. Antigenic and allergenic properties of the storage mite Tyrophagus putrescentiae. Journal of Allergy and Clinical Immunology. 1984/08/01/ 1984;74(2):166–171. doi:https://doi.org/10.1016/0091-6749(84)90281-1

9. Arlian LG, Vyszenski-Moher DL, Johansson SGO, van Hage-Hamsten M. Allergenic Characterization of Tyrophagus putrescentiae Using Sera from Occupationally Exposed Farmers. Annals of Allergy, Asthma & Immunology. 1997/12/01/ 1997;79(6):525–529. doi:https://doi.org/10.1016/S1081-1206(10)63060-8

10. Mullen GR, Oconnor BM. Chapter 26 - Mites (Acari). In: Mullen GR, Durden LA, eds. Medical and Veterinary Entomology (Third Edition). Academic Press; 2019:533–602.

11. Jeong KY, Park JW, Hong CS. House dust mite allergy in Korea: the most important inhalant allergen in current and future. Allergy Asthma Immunol Res. Nov 2012;4(6):313–25. doi:10.4168/aair.2012.4.6.313

12. Zhang C, Li J, Lai X, et al. House dust mite and storage mite IgE reactivity in allergic patients from Guangzhou, China. Asian Pacific journal of allergy and immunology. Dec 2012;30(4):294–300.

13. Choi B, Lee Y, Song T, et al. Sensitization to Tyrophagus putrescentiae in Korean Children with Allergic Diseases. Journal of Allergy and Clinical Immunology. 2010;125(2):AB18. doi:10.1016/j.jaci.2009.12.102

14. Calderon MA, Linneberg A, Kleine-Tebbe J, et al. Respiratory allergy caused by house dust mites: What do we really know? The Journal of allergy and clinical immunology. Jul 2015;136(1):38–48. doi:10.1016/j.jaci.2014.10.012

15. Tham EH, Lee AJ, Bever HV. Aeroallergen sensitization and allergic disease phenotypes in Asia. Asian Pacific journal of allergy and immunology. Sep 2016;34(3):181–189. doi:10.12932/ap0770

16. Xiong Q, Wan ATY, Tsui SK. A Mini-review of the Genomes and Allergens of Mites and Ticks. Curr Protein Pept Sci. 2020;21(2):114–123. doi:10.2174/1389203720666190719150432

17. Saridomichelakis MN, Marsella R, Lee KW, Esch RE, Farmaki R, Koutinas AF. Assessment of cross-reactivity among five species of house dust and storage mites. Vet Dermatol. Apr 2008;19(2):67–76. doi:10.1111/j.1365-3164.2008.00654.x

18. Gafvelin G, Johansson E, Lundin A, et al. Cross-reactivity studies of a new group 2 allergen from the dust mite Glycyphagus domesticus, Gly d 2, and group 2 allergens from Dermatophagoides pteronyssinus, Lepidoglyphus destructor, and Tyrophagus putrescentiae with recombinant allergens. The Journal of allergy and clinical immunology. Mar 2001;107(3):511–8. doi:10.1067/mai.2001.112264

19. Kim CR, Jeong KY, Yi MH, Kim HP, Shin HJ, Yong TS. Cross-reactivity between group-5 and -21 mite allergens from Dermatophagoides farinae, Tyrophagus putrescentiae and Blomia tropicalis. Mol Med Rep. Oct 2015;12(4):5467–74. doi:10.3892/mmr.2015.4093

20. Pomés A, Davies JM, Gadermaier G, et al. WHO/IUIS Allergen Nomenclature: Providing a common language. Molecular immunology. Aug 2018;100:3–13. doi:10.1016/j.molimm.2018.03.003

21. Liu X-Y, Yang KY, Wang M-Q, et al. High-quality assembly of Dermatophagoides pteronyssinus genome and transcriptome reveals a wide range of novel allergens. Journal of Allergy and Clinical Immunology. 2018/06/01/ 2018;141(6):2268-2271.e8. doi:https://doi.org/10.1016/j.jaci.2017.11.038

22. Gafvelin G, Johansson E, Lundin A, et al. Cross-reactivity studies of a new group 2 allergen from the dust mite Glycyphagus domesticus, Gly d 2, and group 2 allergens from Dermatophagoides pteronyssinus, Lepidoglyphus destructor, and Tyrophagus putrescentiae with recombinant allergens. Journal of Allergy and Clinical Immunology. 2001;107(3):511–518. doi:10.1067/mai.2001.112264

23. Cuevas M, Polk M-L, Becker S, et al. Rhinitis allergica in storage mite allergy. Allergo Journal International. 2022/05/01 2022;31(3):59–68. doi:10.1007/s40629-022-00205-w

24. Reginald K, Tan CL, Chen S, Yuen L, Goh SY, Chew FT. Characterization of Der f 22 - a paralogue of the major allergen Der f 2. Sci Rep. Aug 6 2018;8(1):11743. doi:10.1038/s41598-018-30224-z

25. Fujimura T, Aki T, Isobe T, et al. Der f 35: An MD-2-like house dust mite allergen that cross-reacts with Der f 2 and Pso o 2. Allergy. Nov 2017;72(11):1728–1736. doi:10.1111/all.13192

26. Zhou Y, Li L, Yu Z, et al. Dermatophagoides pteronyssinus allergen Der p 22: Cloning, expression, IgE-binding in asthmatic children, and immunogenicity. Pediatr Allergy Immunol. Aug 2022;33(8):e13835. doi:10.1111/pai.13835

27. Morales M, Iraola V, Leonor JR, Carnés J. Enzymatic activity of allergenic house dust and storage mite extracts. J Med Entomol. Jan 2013;50(1):147–54. doi:10.1603/me12154

28. Song H, Lee J, Jeong KY, Cheon D-S, Park J-W. Comparison of sensitization pattern to dust mite allergens between atopic dermatitis patients and dogs and non-specific reactivity of canine IgE to storage mite, Tyrophagus putrescentiae. 2022;

29. Arruda LK, Vailes LD, Platts-Mills TA, et al. Sensitization to Blomia tropicalis in patients with asthma and identification of allergen Blo t 5. Am J Respir Crit Care Med. Jan 1997;155(1):343–50. doi:10.1164/ajrccm.155.1.9001334

30. Gao YF, Wang de Y, Ong TC, Tay SL, Yap KH, Chew FT. Identification and characterization of a novel allergen from Blomia tropicalis: Blo t 21. The Journal of allergy and clinical immunology. Jul 2007;120(1):105–12. doi:10.1016/j.jaci.2007.02.032

31. Park JW, Ko SH, Yong T-S, Ree H-I, Jeoung B-J, Hong C-S. Cross-reactivity of Tyrophagus putrescentiae with Dermatophagoides farinae and Dermatophagoides pteronyssinus in urban areas. Annals of Allergy, Asthma & Immunology. 1999;83(6):533–539.

32. Barber D, Arias J, Boquete M, et al. Analysis of mite allergic patients in a diverse territory by improved diagnostic tools. Clin Exp Allergy. Jul 2012;42(7):1129–38. doi:10.1111/j.1365-2222.2012.03993.x

33. Huang CH, Liew LM, Mah KW, Kuo IC, Lee BW, Chua KY. Characterization of glutathione S-transferase from dust mite, Der p 8 and its immunoglobulin E cross-reactivity with cockroach glutathione S-transferase. Clin Exp Allergy. Mar 2006;36(3):369–76. doi:10.1111/j.1365-2222.2006.02447.x

34. Stewart GA. Studies of house dust mites can now fully embrace the “-omics” era. The Journal of allergy and clinical immunology. Feb 2015;135(2):549–50. doi:10.1016/j.jaci.2014.12.934

35. Chan T-F, Ji K-M, Yim AK-Y, et al. The draft genome, transcriptome, and microbiome of Dermatophagoides farinae reveal a broad spectrum of dust mite allergens. Journal of Allergy and Clinical Immunology. 2015/02/01/ 2015;135(2):539–548. doi:https://doi.org/10.1016/j.jaci.2014.09.031

