## Supplementary Materials for "Multi-Omic Analysis of *Tyrophagus putrescentiae* Reveals Insights into the Allergen Complexity of Storage Mites"

for

**Contents:**

Methods

Supplementary Tables E1–2

Supplementary Figures E1–8

References

### Methods

*Genome and transcriptome data*

The genome and transcriptome data of *T. putrescentiae* have been reported [1] and deposited under in the NCBI database under BioProject accession PRJNA706095. The GenBank assembly accession of is GCA_021730765.1 and the SRA accession of transcriptome data is SRR13837414 that contains three sets of original reads.

*In silico identification of allergens*

The protein sequences of allergens in astigmatic mites were downloaded from the WHO/IUIS allergen nomenclature database [2] (dated March 2022) as references. The reference allergens were searched in the annotated protein sequences of *T. putrescentiae* genome [1] (GenBank assembly accession: GCA_021730765.1) using BLASTP v2.9.0 [3] with the options '-evalue 1e-6 -max_hsps 1 -max_target_seqs 1 -outfmt 6'. The top matched proteins were candidate allergens. Then, the transcriptome data were mapped to the genome using Hisat2 [4] and the format conversion was performed by SAMtools [5]. The sorted and indexed bam file was visualized in the Integrative Genomics Viewer (IGV) [6] and used for the manual curation of the candidate allergens. The final *in silico* identified allergens were listed in Table 1 and the sequences have been deposited in NCBI GenBank under accessions OP558975-OP559059.

*Quantification of gene expression level*

To quantify the gene expression level of allergens, all the coding sequences (CDS) were collected and indexed using Salmon v0.12.0 [7] with the options '-i index --type quasi -k 31'. Then the TP1 and TP3 transcriptome reads (NCBI SRA accession: SRR13837414) were mapped to the indexed CDS of all allergens using Salmon v0.12.0 [7] and the gene expression levels were represented by the TPM (transcript per million). As for the gene expression of Niemann-Pick proteins type C2 (NPC2) family and glutathione S-transferases (GSTs), the CDS of target genes were used as indexes, respectively.

*Cloning, expression, and ELISA of recombinant proteins*

Protein and CDS sequences of six *T. putrescentiae* proteins were obtained after manual curation in the genome. The codon-optimized CDS sequences were synthesized and subcloned into plasmid vector (pET-32a+), and then expressed into recombinant proteins in TOP10 *E. coli*. The cloning and expression of the recombinant proteins were performed by Sangon Biotech (Shanghai, China).

The allergenicity of the recombinant proteins was assessed by the enzyme-linked immunosorbent assay (ELISA). For the experiments, serum samples were obtained from 18 patients who presented with positive SPT or ImmunoCAP results towards *T. putrescentiae* (d72), and from 13 healthy individuals who served as negative controls (Table 1). In the indirect sandwich ELISA, a 96-well microtiter plate was coated with 100 ul (2 ug/ml) of recombinant proteins per well in sodium bicarbonate coating buffer (0.1 mol/L, pH 9.5). The plate was washed by 0.05% Tween-20 in PBS washing buffer for 4 times and then was blocked with 375 ul 3% skim milk in PBST buffer for 30 minutes at 37°C. Serum samples were diluted 1:4 in PBST containing 1% BSA and 50 ul was added to each blocked well for 1 hr at room temperature. After washing, IgE antibodies were detected using biotinylated goat anti-human IgE (Vector, Burlingame, CA, USA) and streptavidin peroxidase (Sigma-Aldrich, St. Louis, MO, USA). The wells were then washed again with PBST buffer and incubated with 100 ul 3,3',5,5'-Tetramethylbenzidine (Kirkegaard and Perry Laboratories, Gaithersburg, MD, USA) for 15 mins at room temperature. The reactions were ceased with the addition of 2M sulphuric acid (50 ul/well). The plates were then placed in the Benchmark Plus Microplate Spectrophotometer (BioRad, Hercules, CA, USA) and absorbance at 450 nm was recorded, where a high absorbance is indicative of a high concentration.

*2D gel electrophoresis, immunoblotting and MALDI-TOF mass spectrometry*

Two-dimension (2D) gel electrophoresis was performed by Shenzhen University. Serum samples were obtained at the First Affiliated Hospital of The First Affiliated Hospital of Guangzhou Medical University from 25 patients (mean age, 19.9 years; range, 2–51 years, Table 2) who presented with allergic rhinitis and/or asthma and were examined for dust mite allergies.

In brief, the final concentration of total protein extracted from *T. putrescentiae* was adjusted to 1.5 ug/ul. 2D- polyacrylamide gel electrophoresis (PAGE) was performed in duplicate using 300 ug total protein per gel. The first gel was used for IgE-blot analysis to locate dust mite antigens, while the second gel was stained with Coomassie blue (Sigma, St. Louis, MO, USA) for cutting spots of allergen candidates. These putative allergen spots were then processed to MALDI-TOF mass spectrometry analysis.

In 2D gel electrophoresis, the first-dimension separation step was to separate protein mixture by the isoelectric point (pI). Isoelectric focusing (IEF) was performed by two immobilized pH gradient

(IPG) strips (pH 3.0–10.0 and pH 4.0–7.0). In each gel, 300 ug proteins were used and make up to 125 ul using 8M Urea, 2% CHAPS, 65 mM DTT, 0.5% v/v IPG buffer pH 3-10 and 0.001% bromophenol blue. The entire 7 cm precast gel tapes were soaked in 125 ul protein extracts for an hour and covered with mineral oil. The focus tray was put into the IEF instrument for running (operating voltage followed by 50 V 12 hrs, 250 V 30 mins, 500 V 30 mins, 4 kV 3 hrs, 4 kV 20,000 Vh). After the isoelectric focusing, the strips were washed with distilled water and then put into the equilibration buffer I (0.375 mol/L Tris-HCl, 6 mol/L urea, 20% glycerol, 2% SDS, 2% DTT) for 15 mins and rinsed with distilled water. The strips were then shaken gently for 15 mins in equilibration buffer II (0.375 mol/L Tris-HCl, 6 mol/L urea, 20% glycerol, 2% SDS, 2.5% iodoacetamide), and rinsed with distilled water.

The second-dimension electrophoresis was used to separate proteins according to their molecular mass in a 12% sodium dodecyl sulfate polyacrylamide gel electrophoresis (SDS-PAGE) gel. Prior to SDS-PAGE, an equilibration step was applied to the first gel. The separated proteins on IEF gel were complexed with SDS, reduced with DTT, and then alkylated with iodoacetamide. After that, the equilibrated strip was loaded horizontally into an SDS-PAGE gel and was secured by overlaying with molten agarose solution. The gel was loaded into the electrophoresis cell and run in 60 V for 20 mins and then 120 V voltage for 2 hrs. The separated proteins were then transferred from the gels onto PVDF membranes. The PVDF membranes were blocked with 3% bovine serum albumin (BSA) at 4 °C overnight, and incubated with pooled sera (dilution ratio, 1:10) from 25 patients, followed by applying an anti-human biotin-conjugated secondary antibody. IgE-allergen bound spots were conjugated with mouse anti-human IgE Horseradish Peroxidase Streptavidin (HRP) (1:2000) and signals were developed with 3,3'-diaminobenzidine (DAB). Spots in another parallel Coomassie blue stained gel were excised according to the corresponding position of the IgE-bound spots. After trypsin digestion, the excised protein samples were analyzed by MALDI-TOF mass spectrometry.

The peptide fingerprinting and bioinformatic identification of the IgE-bound proteins is achieved using MALDI-TOF mass spectrometry. The Coomassie blue-stained protein spots were destained with 50% methanol/10 mM ammonium bicarbonate (NH4HCO3) solution, and then dehydrated repeatedly with acetonitrile (ACN). The gel spots were dried in a vacuum centrifuge and rehydrated with freshly prepared trypsin enzyme in a buffer (20 ng/ul trypsin in 10 mM Ammonium bicarbonate), followed by incubating on ice for 15 mins. The gel spots were incubated overnight at 30 °C for digestion. Half volume of 80% acetonitrile/ 2.5% trifluoroacetic acid (TFA) was added to each tube, and each extract was sonicated for 10 minutes.

Digested samples (0.5 ul each) were spotted onto stainless steel MALDI-plates and leave to dry at room temperature, followed by 0.5 ul matrix solution (5 mg/ml α-Cyano-4-hydroxycinnamic acid in 50% CAN/0.5% TFA). The instrument and MALDI plate were calibrated by a calibration mixture (desArgl-Bradykinin, Angiotensin I, Glul-Fibrinopeptide B and Neurotensin). The target plate was inserted into ABI 4700 TOF-TOF Proteomics Analyzer (Applied Biosystems, Framingham, MA, USA) instrument. All the mass spectrometry spectra were acquired using reflectron positive-ion mode over mass range m/z 700-4000 with acceleration at 20 kV in batch mode. Five to eight abundant peptide precursor ions were chosen automatically for tandem MS/MS analysis with collision-induced dissociation (CID) for each sample. Air was used as collision gas and the collision energy was 1 kV.

The annotated protein sequences of *T. putrescentiae* were used as the database for peptide search. The mass spectrometry data were searched by GPS Explorer using the search engine MASCOT (Matrix Sciences, London, United Kingdom). For peptide fingerprinting, the searches were performed with a peptide mass tolerance of 50 ppm. Significant hits were determined by Mascot probability analysis and target with at least two matches were accepted. Combined (MS+MS/MS) analysis was performed for proteins failed to be identified by peptide fingerprinting. The search was performed with a peptide mass tolerance of 50 ppm, MS/MSion mass tolerance of 0.1 Da, one missed cleavage, and oxidation of methionine.

*Comparative analysis of gene families*

All proteins of the six astigmatic mites were searched by BLASTP v2.9.0 [3] with reference proteins at E-value cutoff of 1E-6. All the group 2, 22 and 35 allergens of astigmatic mite in the WHO/IUIS allergen nomenclature database [2] (dated March 2022) were used for searching NPC2 family genes, while all GSTs in Swiss-Prot database [8] (dated October 2021) were used for the GSTs of mites. After manual curation based on transcriptome data to confirm intron-exon split sites and filtering out annotated proteins with shorter than 50% average length of the gene family, all proteins of the six astigmatic mites identified in the target gene families were aligned and drawn into a phylogenetic tree. Sequence alignment was performed by CLUSTAL W [9] and MUSCLE [10] in MEGA v11.0.13 [11], and all phylogenetic trees were constructed by MEGA v11.0.13 [11] with maximum likelihood (ML) algorithm in the JTT (Jones-Taylor-Thornton) model, 80% site coverage and 100 bootstrap replicates, and then edited by the online tool Interactive Tree of Life (iTOL) [12].

If two genes were located adjacently on genome and no other gene was located between them, they were considered as tandemly arrayed genes. If two genes were separated by no more than 10 genes, they were considered as proximally arrayed. Frequent tandemly arrayed genes were identified in gene families and connected with curve lines in phylogenetic trees edited by the online tool iTOL [12] with validated gene synteny information.

*Ethics approval*

This study was approved by the institutional review board (IRB no. 4-2013-0397) of Institute of Allergy, College of Medicine, Yonsei University for using the patient sera in ELISA experiments and the hospital ethics committee of The First Affiliated Hospital of Guangzhou Medical University (reference no. 2017018) for using the pooled patient sera in immunoblotting (western blotting) following the 2D gel electrophoresis.

### Supplementary Tables

#### Table E1. Metadata of the patient sera for ELISA experiment

Serum samples were collected by Institute of Allergy, Department of Internal Medicine, College of Medicine, Yonsei University (Seoul, Korea). All the patient sera were tested to be positive in SPT or ImmunoCAP towards *T. putrescentiae* (ImmunoCAP allergen d72). In the SPT column, H means histamine. The ELISA results were that of the whole protein extract (Fig. 2F).

| patient sera (1:4 diluted) | SPT (wheal/erythema, cm) | ImmunoCAP | ELISA |
|---|---|---|---|
| **1** | d72 (2*2/2*2) H (5*4/40*25) | -- | - |
| **2** | d72 (2*2/10*10) H (6*6/30*30) | -- | - |
| **3** | d72 (3*3/20*10) H (5*5/25*15) | -- | + |
| **4** | d72 (4*4/20*20) H (5*4/20*20) | -- | - |
| **5** | d72 (2*2/20*20) H (6*5/30*30) | -- | + |
| **6** | d72 (2*2/20*20) H (3*2/30*30) | -- | - |
| **7** | d72 (2*2/30*20) H (5*4/30*25) | -- | + |
| **8** | d72 (5*5/15*15) H (4*4/20*10) | -- | + |
| **9** | d72 (3*3/20*15) H (5*5/15*15) | -- | - |
| **10** | d72 (2*2/10*10) H (3*3/10*10) | -- | + |
| **11** | d72 (4*3/15*10) H (3*3/10*10) | -- | + |
| **12** | d72 (3*3/7*5) H (4*4/15*15) | -- | - |
| **13** | d72 (3*2/10*10) H (4*3/15*15) | -- | + |
| **14** | d72 (3*2/10*10) H (5*4/20*20) | -- | + |
| **15** | d72 (3*3/25*15) H (4*3/20*15) | -- | + |
| **16** | d72 (4*3/20*20) H (4*3/30*20) | -- | - |
| **17** | -- | 4+ | + |
| **18** | -- | 3+ | + |

**Table E2. Metadata of the patient sera pooled for 2D gel electrophoresis**

Serum samples were obtained at The First Affiliated Hospital of Guangzhou Medical University (Guangzhou, China) from 25 patients (mean age, 19.9 years; range, 2–51 years) who presented with allergic rhinitis and/or asthma and were examined to be dust mite allergy.

| Patient | Sample | Sex | Age | Diagnosis | d1 (*D. pteronyssinus*) * | |
|---|---|---|---|---|---|---|
| | | | | | Conc. (kU/l) | Level |
| **85909** | BJ1905 | F | 17yr | Allergic purpura | 16.6 | 3 |
| **85963** | BJ1964 | M | 51yr | Skin allergy | 14.2 | 3 |
| **86332** | BGM8857 | F | 3yr | Rhinitis | 8.7 | 3 |
| **86432** | BGM9094 | F | 2yr | Asthma | 5.63 | 3 |
| **87631** | BJ1085 | M | 33yr | Allergic dermatitis | 17.4 | 3 |
| **87649** | BJ1053 | M | 9yr | Allergic rhinitis | 35.5 | 4 |
| **88216** | BJ2203 | M | 6yr | Acute suppurative tonsillitis | 6.59 | 3 |
| **89070** | BJ3542 | F | 26yr | Rhinitis | 13.8 | 3 |
| **89260** | BGM9501 | F | 42yr | Allergic rhinitis | 38.2 | 4 |
| **89580** | BJ4754 | F | 12yr | Systemic lupus erythematosus | 7.96 | 3 |
| **90444** | BJ6422 | F | 16yr | Allergic purpura | 5.7 | 3 |
| **90996** | BJ7531 | M | 42yr | Acute urticaria | 5.74 | 3 |
| **91503** | BJ8361 | F | 21yr | Allergic dermatitis | 4.33 | 3 |
| **91506** | BJ8353 | F | 18yr | Allergic dermatitis | 3.83 | 3 |
| **91510** | BJ8356 | M | 7yr | Allergic dermatitis | 25 | 4 |
| **91508** | BJ8380 | M | 4yr | Allergic purpura | 26.1 | 4 |
| **91523** | BJ8443 | M | 39yr | Acute urticaria | 14.3 | 3 |
| **91524** | BJ8422 | F | 17yr | Allergic dermatitis | 80.4 | 5 |
| **91525** | BJ8426 | M | 7yr | Asthmatic bronchitis | 85.3 | 5 |
| **91580** | BGM102 | F | 32yr | Urticaria | 11.1 | 3 |
| **91582** | BGM150 | M | 36yr | Urticaria | 4.5 | 3 |
| **91253** | BJ7960 | M | 16yr | Allergic purpura | > 100 | 6 |
| **91554** | BJ8525 | F | 13yr | Urticaria | > 100 | 6 |
| **91555** | BJ8543 | M | 12yr | Urticaria | 72.9 | 5 |
| **91557** | BJ8462 | F | 17yr | Chronic urticaria | 3.93 | 3 |

* These patient sera samples (obtained at The First Affiliated Hospital of Guangzhou Medical College, Guangzhou, China) were tested with ImmunoCAP kit (d1 for *D. pteronyssinus*) and their IgE reactivity were at least level 3 (IgE $\geq$ 3.5 kU/l).

### Supplementary Figures

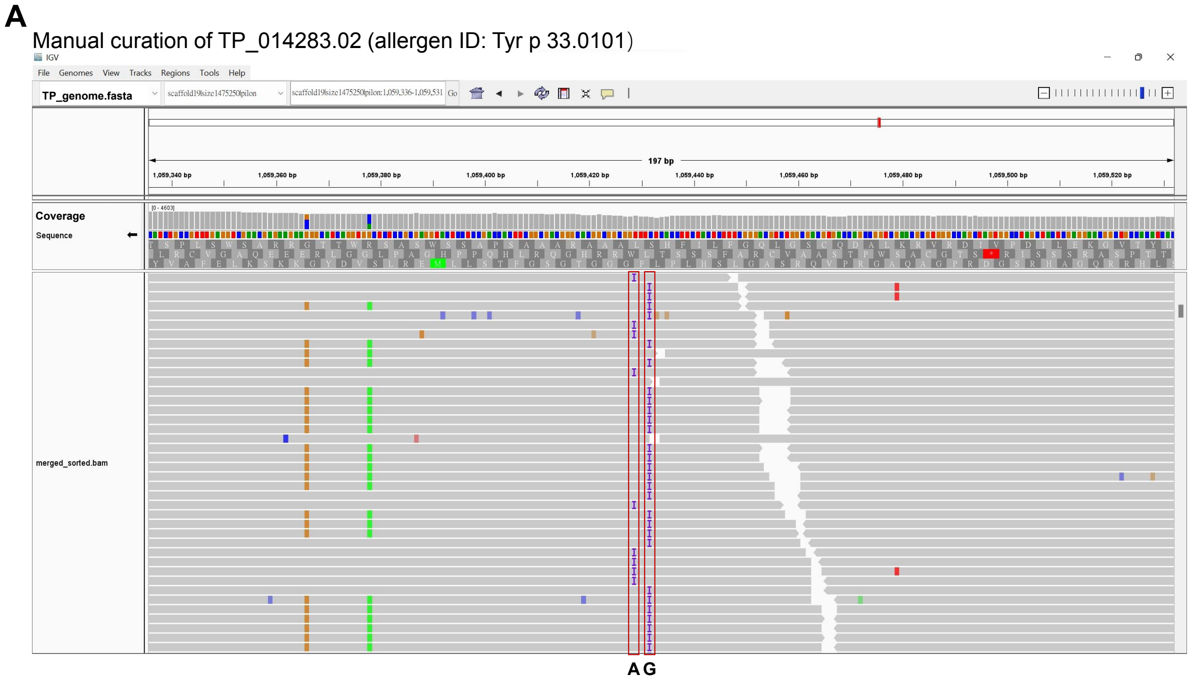

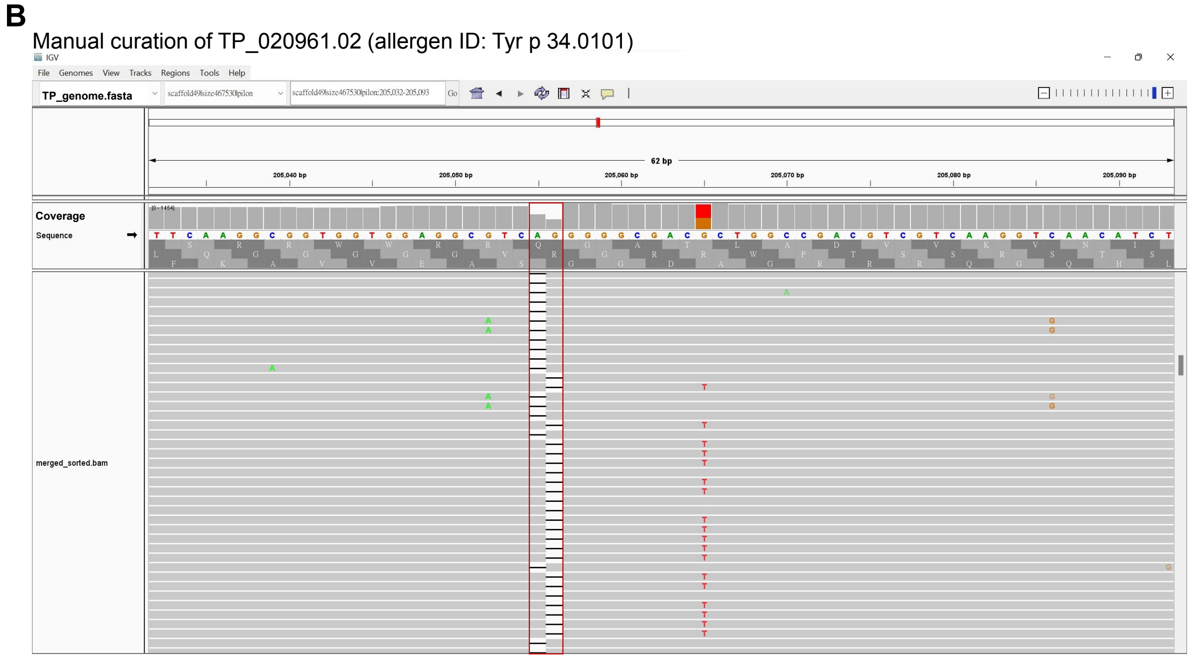

**Fig. E1. Examples of manual curation**

**(A)** Manual curation of TP_014283.02 (allergen ID: Tyr p 33.0101) in Integrative Genomics Viewer (IGV). Two genotypes of false insertions were marked in red squares. **(B)** Manual curation of TP_020961.02 (allergen ID: Tyr p 34.0101). Two genotypes of false deletions were marked in red square.

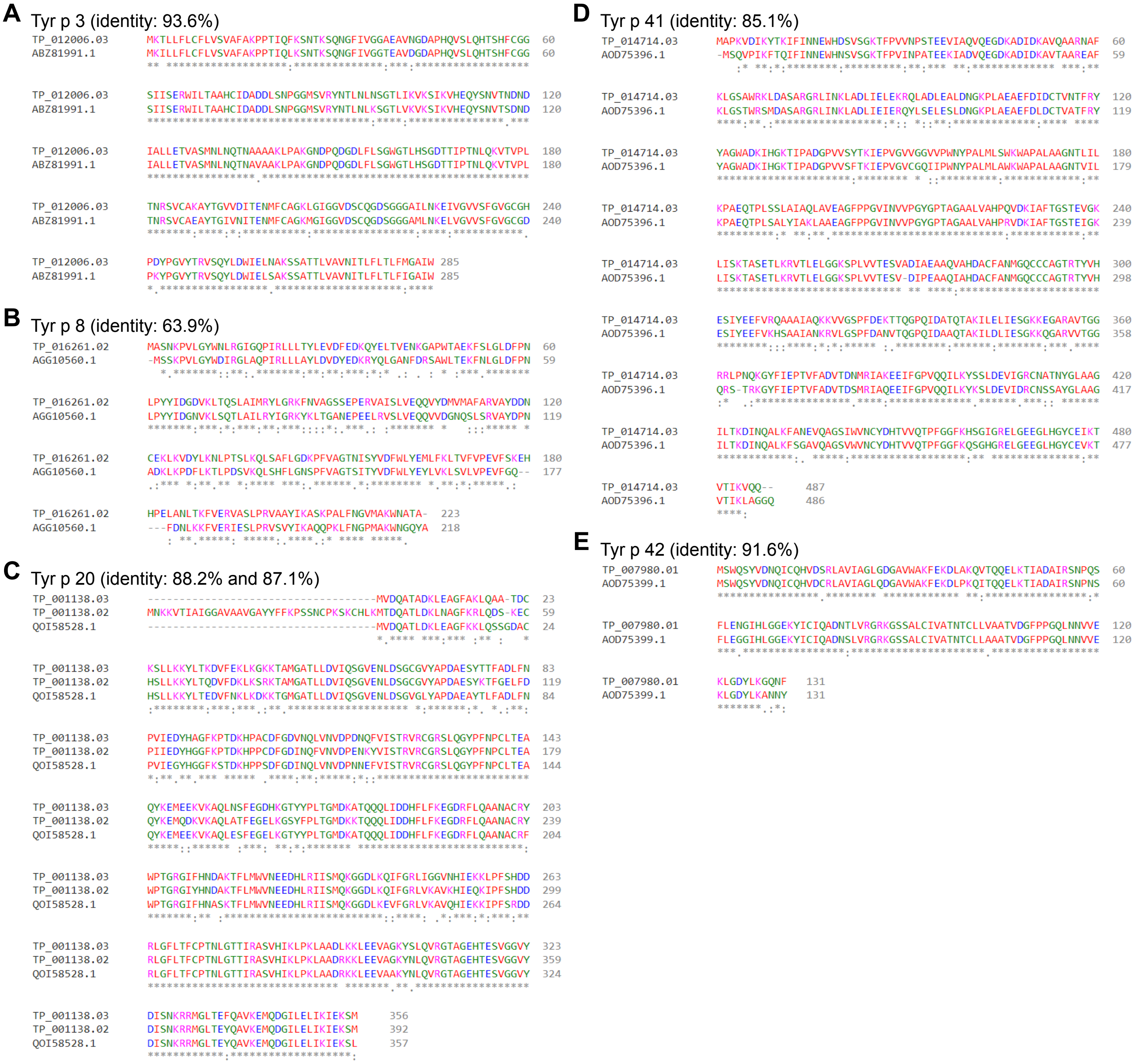

**Fig. E2. Sequence alignment of Tyr p 3, 8, 20, 41 and 42**

**(A)** Sequence alignment of Tyr p 3.0101 (gene locus: TP_012006.03) and the reported Tyr p 3 (GenBank accession: ABZ81991.1). **(B)** Sequence alignment of Tyr p 8.0101 (gene locus: TP_016261.02) and the reported Tyr p 8 (GenBank accession: AGG10560.1). **(C)** Sequence alignment of Tyr p 20.0101 (gene locus: TP_001138.03), Tyr p 20.0201 (gene locus: TP_001138.02) and the reported Tyr p 20 (GenBank accession: QOI58528.1). **(D)** Sequence alignment of Tyr p 41.0101 (gene locus: TP_014714.03) and the reported Tyr p 41 (GenBank accession: AOD75396.1). **(E)** Sequence alignment of Tyr p 42.0101 (gene locus: TP_007980.01) and the reported Tyr p 42 (GenBank accession: AOD75399.1). All the sequence alignments were performed by the online tool Clustal Omega.

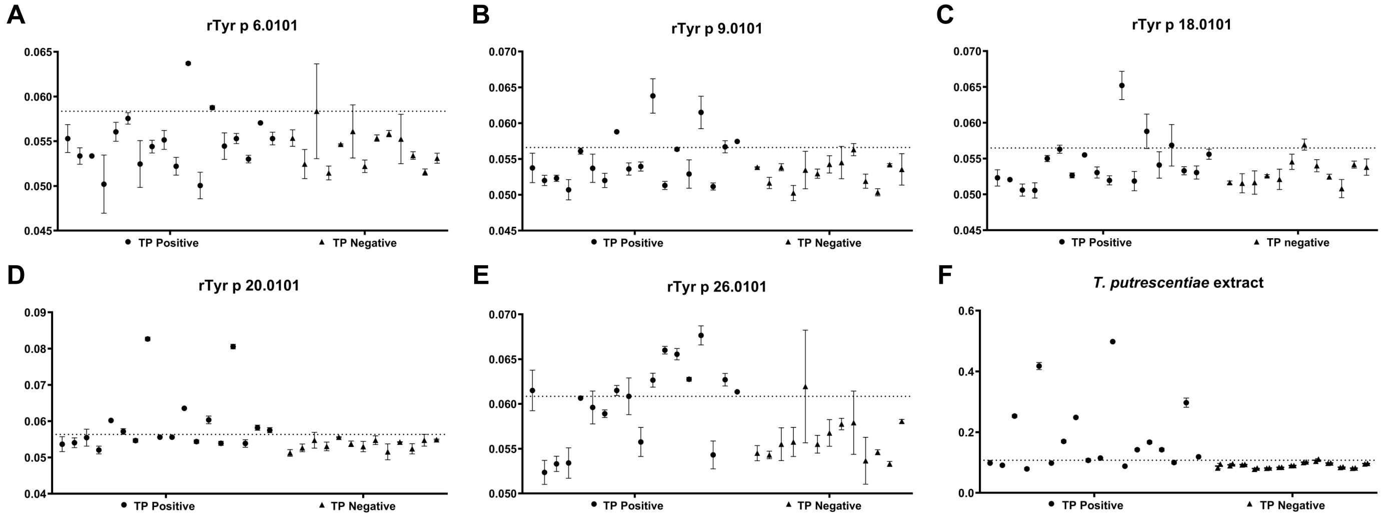

**Fig. E3. ELISA results for recombinant proteins and crude protein extract**

The allergenicity of recombinant proteins and whole protein extract of *T. putrescentiae* was tested by ELISA using sera samples from 18 allergy patients (TP positive) and 13 healthy individuals (TP negative) as negative controls (Table E1). The dotted lines indicate the cutoff value, mean+ 2*SD (standard deviation) of the negative controls. The allergenicity of recombinant proteins was as follows: **(A)** rTyr p 6.0101 (11.1%), **(B)** rTyr p 9.0101 (22.2%), **(C)** rTyr p 18.0101 (11.1%), **(D)** rTyr p 20.0101 (44.4%), **(E)** rTyr p 26.0101 (50.0%). Also, 11/18 (61.1%) of the patient serum samples showed positive responses to the crude protein extract **(F)**.

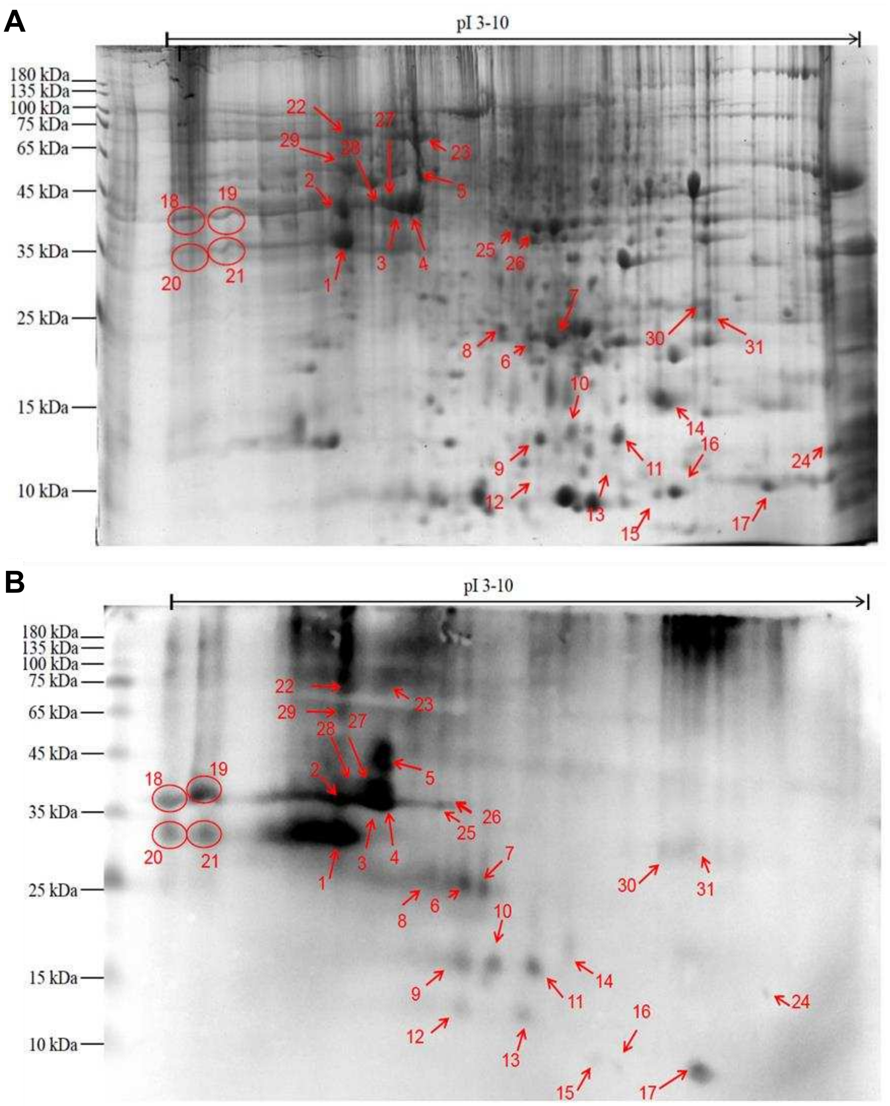

**Fig. E4.** ***T. putrescentiae*** **proteins in two-dimensional (2D) gel electrophoresis**
**(A)** SDS-PAGE image. The Coomassie blue stained 2D gel showed that the success of the separation of more than 60 *T. putrescentiae* proteins. **(B)** Western blotting image. The results of immunoblotting from the 2D gel showed that there were 31 protein spots in the total protein of *T. putrescentiae* bound by the specific IgE in the patient sera (Table E2), numbered 1-31. All the protein spots could be found correspondingly in the 2D gel.

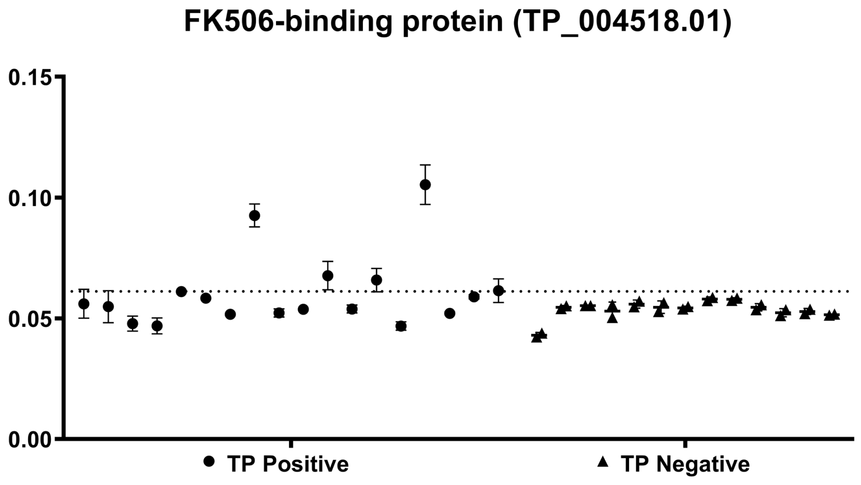

**Fig. E5. ELISA result for the recombinant FK506-binding protein (TP_004518.01)**
The allergenicity of the recombinant FK506-binding protein (TP_004518.01) of *T. putrescentiae* was tested by ELISA using sera samples from 18 allergy patients (TP positive) and 13 healthy individuals (TP negative) as negative controls (Table E1). The dotted lines indicate the cutoff value, mean+ 2*SD (standard deviation) of the negative controls. The allergenicity 22.2% (4/18).

```
CLUSTAL O(1.2.4) multiple sequence alignment

DF_001154.01      MISKILCLSLLV--------------------------------------AAVVADQVDV  22
DP_010248.01      MMYKILCLSLLV--------------------------------------AAVAADQVDV  22
Eurm_2.0101       -MYKILCLSLLV--------------------------------------AAVAADQVDI  21
SS_011027.01      MLKYFLASVLLVAAYSLVISAYDQNDEFDQHSTSKSKQPETITTNKPDAKNPKDFRDIAF  60
                   :  :*.  ***                                            :: .

DF_001154.01      KDCANNEIKKVMVDGCHGSDPCIIHRGKPFTLEALFDANQNTKTAKIEIKASL-DGLEID  81
DP_010248.01      KDCANHEIKKVLVPGCHGSEPCIIHRGKPFQLEAVFEANQNSKTAKIEIKASI-DGLEVD  81
Eurm_2.0101       KDCANHEIKKVMVPGCKGSEPCVIHRGTAFQLEAVFDANQNSNAAKIEIKATI-DGVEID  80
SS_011027.01      TDCGNHELLHVRLTGCRGSIPCVLYRKSVVHLTAIFKSNQNTTTAVVGLQATLPGGLELP  120
                  .**.*:*: :* : **:** **:::* . . * *:*.:***:.:* : ::*:: .*:*:

DF_001154.01      VPGIDTNACHFVKCPLVKGQQYDIKYTWNVPKIAPKSENVVVTVKLIGDNGVLACAIATH  141
DP_010248.01      VPGIDPNACHYMKCPLVKGQQYDIKYTWNVPKIAPKSENVVVTVKVMGDNGVLACAIATH  141
Eurm_2.0101       VPGIDNNLCHFMKCPLVKGQEYDIKYTWNVPRIAPKSENVVVTVKLLGDNGVLACAIATH  140
SS_011027.01      VPGVDTNACHHEHCPITCGSLVTFTYPWVVPKFMPQFSNLTLKAYMEGEHGRMACGIITN  180
                  ***:* * **. :**:. *.   :.* * **:: *: .*:.:.. : *::* :**.* *:

DF_001154.01      AKIRD 146
DP_010248.01      AKIRD 146
Eurm_2.0101       AKIRD 145
SS_011027.01      VSIR- 184
                  ..**
```

**Fig. E6. Multiple sequence alignment of N2 Cluster**

Protein sequences of DF_001154.01, DP_010248.01, Eur m 2.0101 and SS_011027.01 were aligned by the online tool Clustal Omega. A fragment of unaligned sequences of SS_011027.01 was marked in the black square.

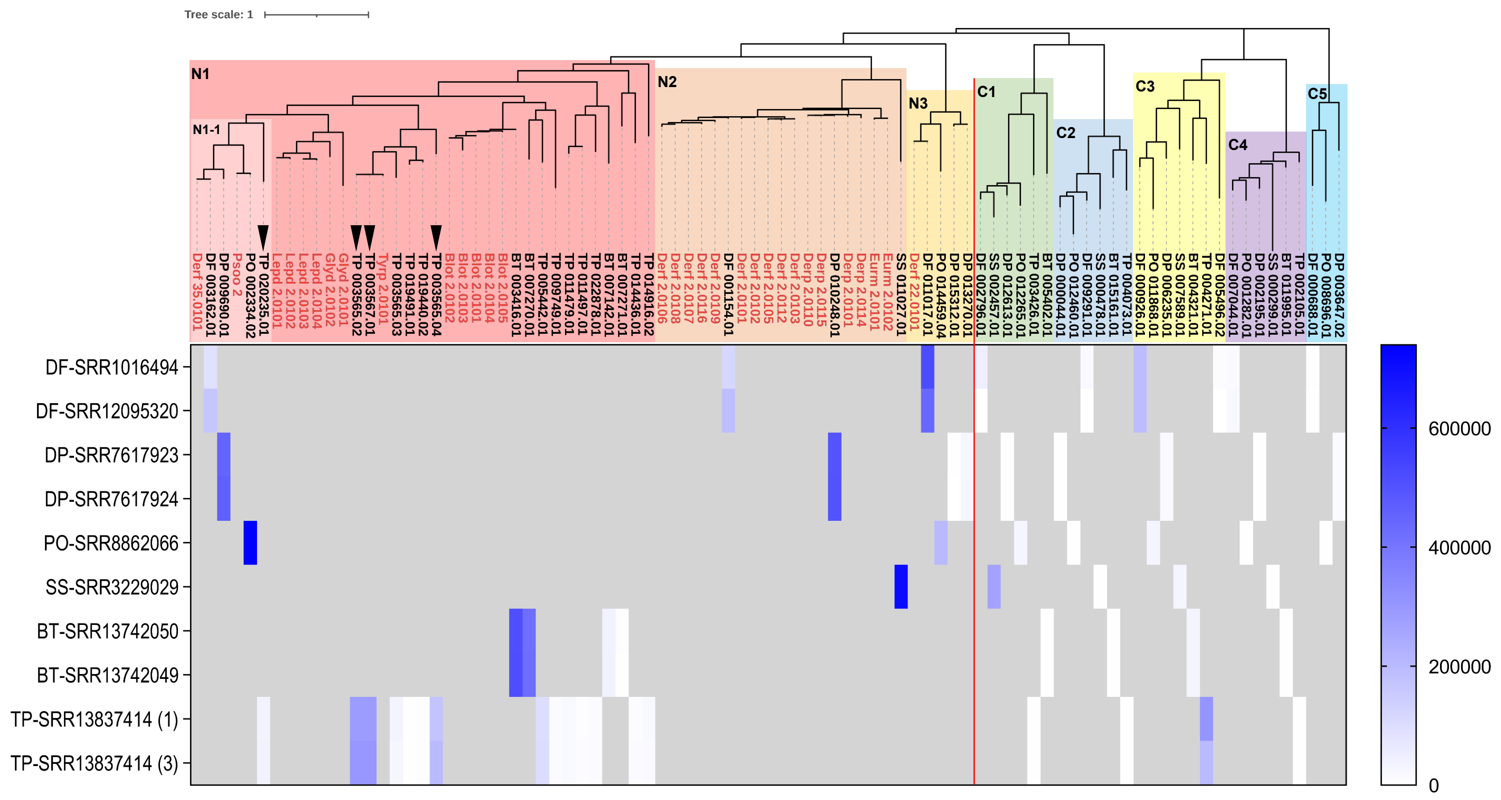

**Fig. E7. Expression levels of NPC2 family**

Gene expression levels of NPC2 family of six astigmatic mites were quantified as TPM using transcriptome data of adult mites. The SRA accessions were labeled in the axis. TP1 and TP3 of SRR13837414 transcriptome data were used for genes of *T. putrescentiae* (labeled 1 and 3, respectively).

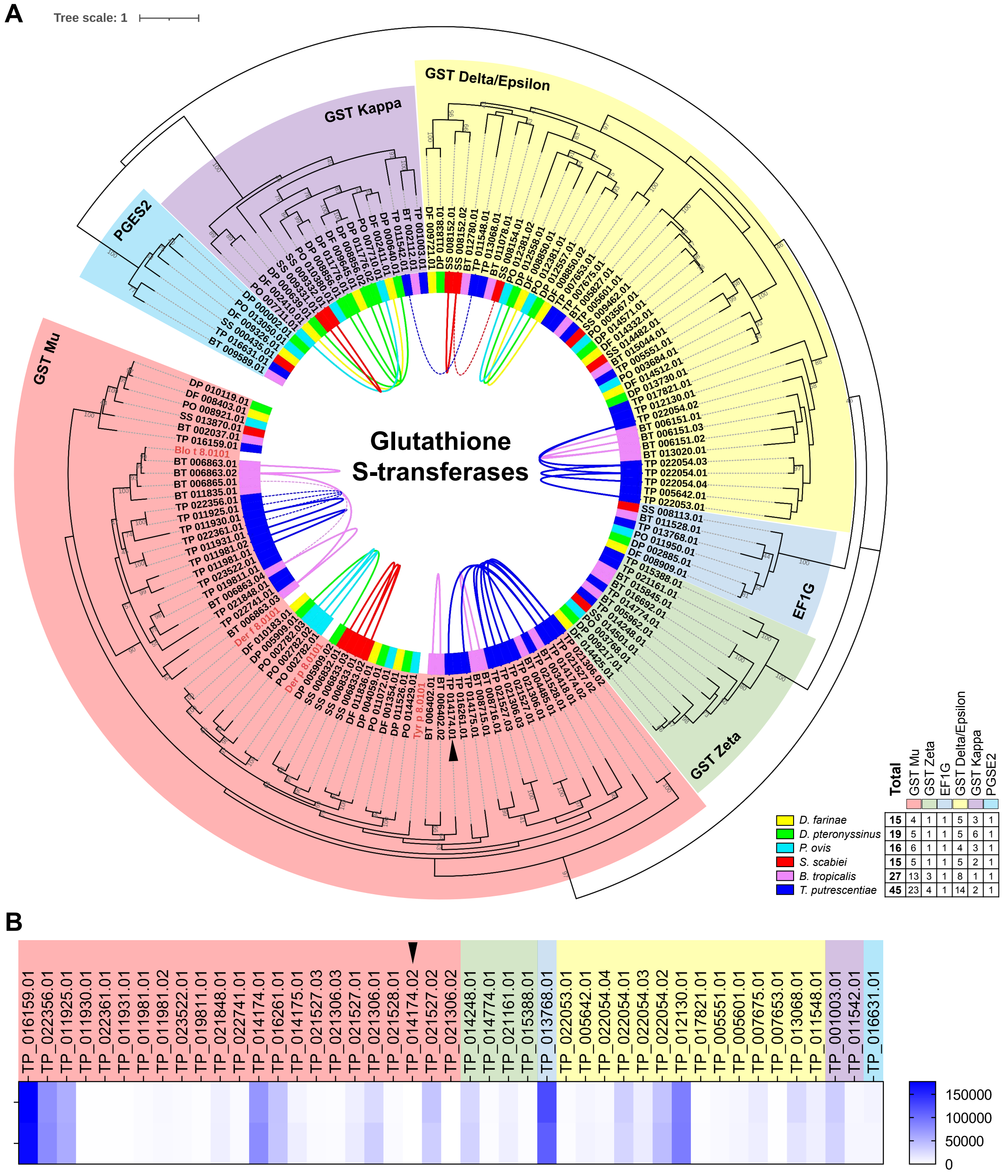

**Fig. E8. Phylogenetic analysis and expression levels of GSTs**

**(A)** Phylogenetic analysis of GSTs of six astigmatic mites. The GST genes were divided into six clusters, including EF1G that contains a eukaryotic elongation factor 1 gamma (eEF1γ) conserved domain on C terminal and prostaglandin E synthase 2 (PTGES2) that has GST activity. The GST allergens including group 8 of astigmatic mites (highlighted in red) were collected from the WHO/IUIS nomenclature database. Tandemly arrayed genes and proximally arrayed genes (separated by no more than 10 genes) were connected by curved solid lines and dotted lines, respectively. The black triangle marked the GST gene identified in Table 2. **(B)** Gene expression levels of GSTs of *T. putrescentae* were quantified as TPM using two transcriptome data of adult mites (TP1 and TP3 of SRR13837414).

### References

1. Xiong Q, Wan AT-Y, Liu X, et al. Comparative Genomics Reveals Insights into the Divergent Evolution of Astigmatic Mites and Household Pest Adaptations. *Molecular Biology and Evolution*. 2022;39(5)doi:10.1093/molbev/msac097

2. Pomés A, Davies JM, Gadermaier G, et al. WHO/IUIS Allergen Nomenclature: Providing a common language. *Molecular immunology*. Aug 2018;100:3-13. doi:10.1016/j.molimm.2018.03.003

3. Camacho C, Coulouris G, Avagyan V, et al. BLAST+: architecture and applications. *BMC Bioinformatics*. 2009/12/15 2009;10(1):421. doi:10.1186/1471-2105-10-421

4. Kim D, Langmead B, Salzberg SL. HISAT: a fast spliced aligner with low memory requirements. *Nature methods*. 2015/04/01 2015;12(4):357-360. doi:10.1038/nmeth.3317

5. Li H, Handsaker B, Wysoker A, et al. The Sequence Alignment/Map format and SAMtools. *Bioinformatics*. 2009;25(16):2078-2079. doi:10.1093/bioinformatics/btp352

6. Robinson JT, Thorvaldsdóttir H, Winckler W, et al. Integrative genomics viewer. *Nature biotechnology*. Jan 2011;29(1):24-6. doi:10.1038/nbt.1754

7. Patro R, Duggal G, Love MI, Irizarry RA, Kingsford C. Salmon provides fast and bias-aware quantification of transcript expression. *Nature methods*. 2017/04/01 2017;14(4):417-419. doi:10.1038/nmeth.4197

8. Bairoch A, Apweiler R. The SWISS-PROT protein sequence database and its supplement TrEMBL in 2000. *Nucleic acids research*. Jan 1 2000;28(1):45-8. doi:10.1093/nar/28.1.45

9. Thompson JD, Higgins DG, Gibson TJ. CLUSTAL W: improving the sensitivity of progressive multiple sequence alignment through sequence weighting, position-specific gap penalties and weight matrix choice. *Nucleic acids research*. Nov 11 1994;22(22):4673-80. doi:10.1093/nar/22.22.4673

10. Edgar RC. MUSCLE: multiple sequence alignment with high accuracy and high throughput. *Nucleic acids research*. 2004;32(5):1792-7. doi:10.1093/nar/gkh340

11. Tamura K, Stecher G, Kumar S. MEGA11: Molecular Evolutionary Genetics Analysis Version 11. *Mol Biol Evol*. Jun 25 2021;38(7):3022-3027. doi:10.1093/molbev/msab120

12. Letunic I, Bork P. Interactive Tree Of Life (iTOL) v5: an online tool for phylogenetic tree display and annotation. *Nucleic acids research*. 2021;49(W1):W293-W296. doi:10.1093/nar/gkab301
